# Mitotic exit is controlled during anaphase by an Aurora B-Cyclin B1/Cdk1 crosstalk

**DOI:** 10.1101/606517

**Authors:** Olga Afonso, Liam P. Cheeseman, Luísa T. Ferreira, Eurico Morais-de-Sá, Helder Maiato

## Abstract

According to the prevailing “clock” model, chromosome decondensation and nuclear envelope reassembly during mitotic exit are byproducts of Cdk1 inactivation at the metaphase-anaphase transition, controlled by the spindle assembly checkpoint. However, mitotic exit was recently shown to be a function of chromosome separation during anaphase, assisted by a midzone Aurora B phosphorylation gradient - the “ruler” model. Here we reconciled both models by showing that Cyclin B1 degradation continues during anaphase in *Drosophila*, mouse and human cells, including primary tissues. This required APC/C^Cdh1^ activity, and failure to degrade Cyclin B1 during anaphase prevented mitotic exit in a Cdk1-dependent manner. Cyclin B1 localization and half-life during anaphase depended on kinesin-6, which targets Aurora B to the spindle midzone. Mechanistically, we show that anaphase duration is regulated by Aurora B-mediated phosphorylation of Cyclin B1. We propose that a crosstalk between molecular “rulers” and “clocks” licenses mitotic exit only after proper chromosome separation.

## Introduction

The decision to enter and exit mitosis is critical for genome stability and the control of tissue homeostasis, perturbation of which has been linked to cancer (Evan and Vousden, 2001; Hanahan and Weinberg, 2011). While the key universal principles that drive eukaryotic cells into mitosis are well established (Domingo-Sananes et al., 2011; Lindqvist et al., 2009; Rieder, 2011), the mechanistic framework that determines mitotic exit remains ill-defined. The prevailing “clock” model conceives that mitotic exit results from APC/C^Cdc20^-mediated Cyclin B1 degradation as cells enter anaphase, under control of the spindle assembly checkpoint (SAC) (Musacchio, 2015). This allows the dephosphorylation of Cdk1 substrates by PP1/PP2A phosphatases, setting the time for chromosome decondensation and nuclear envelope reformation (NER) (Wurzenberger and Gerlich, 2011), two hallmarks of mitotic exit. This model is supported by the observation that inhibition of PP1/PP2A or expression of non-degradable Cyclin B1 (and B3 in *Drosophila*) mutants arrested cells in anaphase (Afonso et al., 2014; Parry and O’Farrell, 2001; Schmitz et al., 2010; Sigrist et al., 1995; Vagnarelli et al., 2011; Wheatley et al., 1997; Wolf et al., 2006). However, because Cyclin B1 is normally degraded at the metaphase-anaphase transition (Clute and Pines, 1999; Huang and Raff, 1999), the data from non-degradable Cyclin B1 expression could be interpreted as a spurious gain of function due to the artificially high levels of Cyclin B1 that preserve Cdk1 activity during anaphase. Whether Cyclin B1/Cdk1 normally plays a role in the control of anaphase duration and mitotic exit remains unknown.

We have recently uncovered a spatial control mechanism that operates after SAC satisfaction to delay chromosome decondensation and NER in response to incompletely separated chromosomes during anaphase in *Drosophila* and human cells (Afonso et al., 2014; Maiato et al., 2014). The central player in this mechanism is a constitutive midzone-based Aurora B phosphorylation gradient that monitors the position of chromosomes along the spindle axis during anaphase (Afonso et al., 2014; Maiato et al., 2014). Thus, mitotic exit in metazoans cannot simply be explained by a “clock” that starts ticking at the metaphase-anaphase transition, but must also respond to spatial cues as cells progress through anaphase. The main conceptual implication of this “ruler” model is that mitotic exit is determined during anaphase, and not at the metaphase-anaphase transition under SAC control. In this case, a molecular “ruler” that prevents precocious chromosome decondensation and NER would allow that all separated sister chromatids end up in two individualized daughter nuclei during a normal mitosis. Moreover, it provides an opportunity for the correction and reintegration of lagging chromosomes that may arise due to deficient interchromosomal compaction in anaphase (Fonseca et al., 2019) or erroneous kinetochore-microtubule attachments that are “invisible” to the SAC (e.g. merotelic attachments) (Gregan et al., 2011). Interestingly, Aurora B association with the spindle midzone depends on the kinesin-6/Mklp2/Subito (Cesario et al., 2006; Gruneberg et al., 2004) and is negatively regulated by Cdk1 (Hummer and Mayer, 2009). Thus, the establishment of a midzone-based Aurora B “ruler” in anaphase is determined by the sudden drop of Cdk1 activity (the “clock”) at the metaphase-anaphase transition. In the present work we investigate whether and how molecular “rulers” also regulate the “clocks” during anaphase to coordinate mitotic exit in space and time in metazoans.

## Results

### Cyclin B1 continues to be degraded during anaphase and its disappearance is a strong predictor of mitotic exit in metazoans

To investigate a possible role of Cdk1 during anaphase, we started by monitoring Cyclin B1-GFP by spinning-disc confocal microscopy in live *Drosophila* and human cells in culture. Mild induction of Cyclin B1-GFP expression in S2 cells reproduced the localization of endogenous Cyclin B1 in the cytoplasm, mitotic spindle, kinetochores and centrosomes (Bentley et al., 2007; Clute and Pines, 1999; Huang and Raff, 1999; Pines and Hunter, 1991), without altering normal anaphase duration or increasing chromosome missegregation (Figure 1a and Figure 1 – figure supplement 1a, a’ and figure supplement 2a, e, f). In agreement with previous reports (Clute and Pines, 1999; Huang and Raff, 1999), cytoplasmic Cyclin B1-GFP levels decreased abruptly at the metaphase-anaphase transition. However, in contrast to what has been observed in *Drosophila* embryos (Huang and Raff, 1999), the centrosomal pool of Cyclin B1 in S2 cells was the most resistant to degradation and persisted well after anaphase onset, becoming undetectable only ~1 min before DNA decondensation (Figure 1a, c and Figure 1 – figure supplement 2a-c and Video 1). Indeed, complete Cyclin B1 disappearance from centrosomes during anaphase strongly correlated with mitotic exit (Figure 1 – figure supplement 2d).

**Figure 1.**
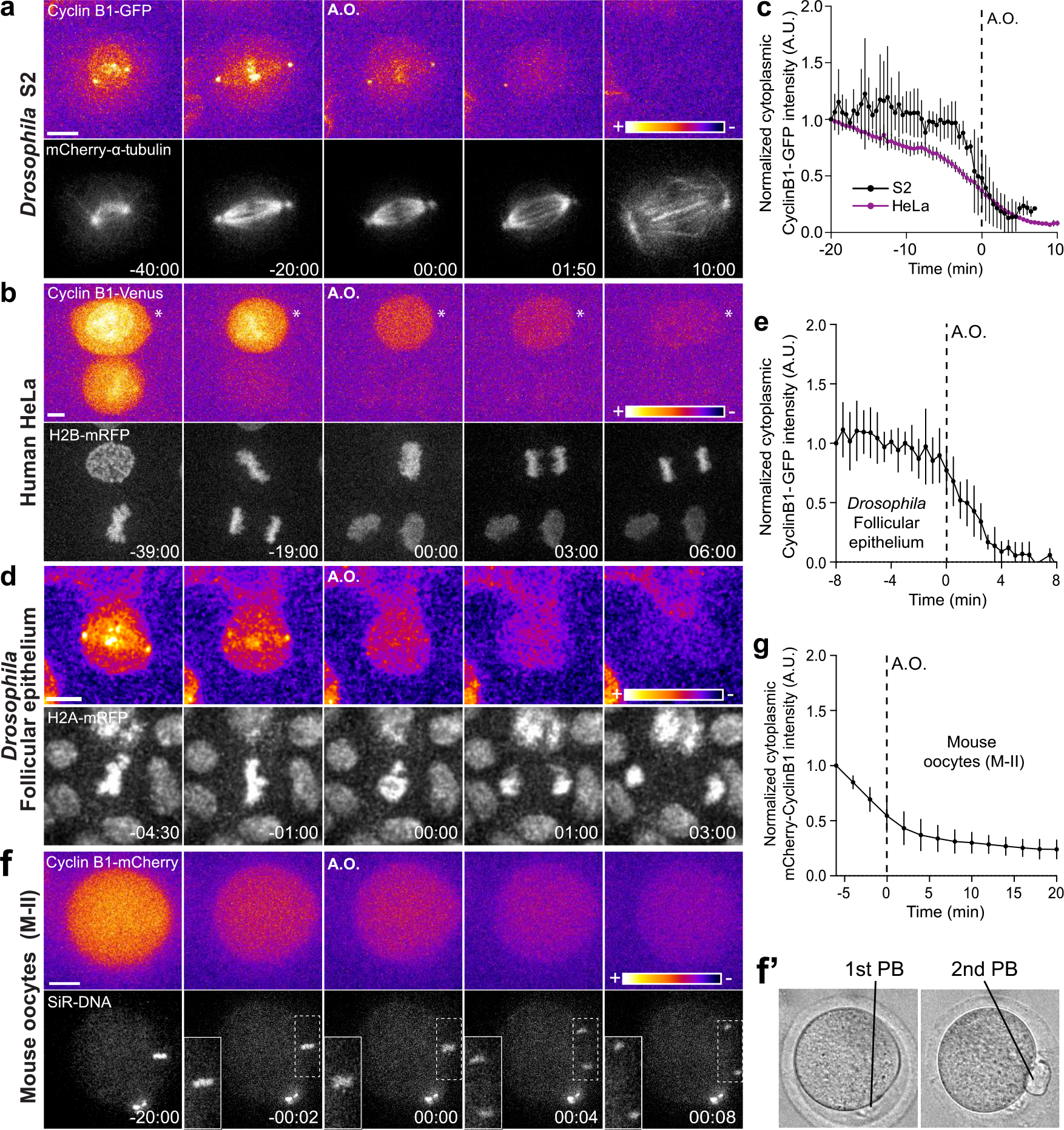
Cyclin B1 continues to be degraded during anaphase. **(a)** *Drosophila* S2 cell from nuclear envelope breakdown to NER showing the different pools of Cyclin B1 in the mitotic apparatus. Scale bar is 5 μm. **(b)** Two neighbor HeLa cells (* indicates a cell that is slightly delayed relative to its neighbor; compare relative Cyclin B1 levels between neighbors as they exited mitosis) expressing exogenous H2B-mRFP and showing continuous degradation of endogenous Cyclin B1-Venus during anaphase. Scale bar is 5 μm. **(c)** Cyclin B1 degradation profile in *Drosophila* S2 cells (n=4 cells) and in HeLa cells with endogenously tagged Cyclin B1-Venus (n=3 cells). Fluorescence intensity values were normalized to 20 min before anaphase onset (A.O.). **(d)** Time-lapse images of dividing *Drosophila* follicle cells expressing endogenously tagged Cyclin B1-GFP and His2Av-mRFP. Scale bar is 5 μm. **(e)** Quantification of Cyclin B1-GFP fluorescence intensity in the cytoplasm (n=12 cells, 7 egg chambers). Fluorescence intensity values were normalized to 8 min before A.O. **(f)** Time-lapse images of a metaphase II oocyte expressing Cyclin B1-mCherry and stained with SiR-DNA undergoing anaphase II after parthenogenic activation. Inset is 1.5× magnification of separating chromosomes. (**f’)** Images of transmission light microscopy showing the same oocyte prior to and after imaging. Note the presence of the first and second polar bodies. Scale bar is 20 μm. **(g)** Quantification of Cyclin B1-mCherry fluorescence intensity in the cytoplasm (n=20 oocytes, 2 independent experiments). Fluorescence intensity values were normalized to 6 min before anaphase onset. The LUT “fire” is used to highlight Cyclin B1 localization in the different systems. Time in all panels is in min:sec.

To extend the significance of these observations, we monitored endogenously tagged Cyclin B1-Venus levels in human HeLa and hTERT-RPE1 cells (Collin et al., 2013) throughout mitosis and found that it is also continuously degraded during anaphase (Figure 1b, c and Figure 1 – figure supplement 3a, c and Videos 2 and 3). The physiological relevance of these findings was confirmed in primary adult *Drosophila* follicular epithelium cells expressing endogenously tagged Cyclin B1-GFP, and after injection of Cyclin B1-mCherry mRNA in mouse oocytes undergoing meiosis II, when chromosomal division is similar to mitosis (Figure 1d-g and Video 4). Thus, Cyclin B1 continues to be degraded during anaphase in *Drosophila*, mouse and human cells, including primary tissues, and its disappearance is a strong predictor of mitotic exit.

### Cyclin B1 degradation and Cdk1 inactivation during anaphase are rate limiting for mitotic exit

To investigate the functional relevance of an anaphase Cyclin B1 pool we acutely inhibited Cdk1 activity at anaphase onset and found that it accelerated NER in S2 cells (Figure 2a-d). In contrast, expression of non-degradable Cyclin B1 induced a delay in anaphase for several hours, as shown previously in human cells and *Drosophila* embryos (Afonso et al., 2014; Parry and O’Farrell, 2001; Potapova et al., 2006; Sigrist et al., 1995; Wolf et al., 2006), and this delay was dependent on Cdk1 activity (Figure 2 - figure supplement 1a-b). However, because expression of non-degradable Cyclin B1 does not reflect a physiological role for Cyclin B1 degradation in mitotic exit, we also inhibited proteasome-mediated degradation of wild-type Cyclin B1 by adding MG132 just before or at anaphase onset in *Drosophila* and human cells, respectively. Under these conditions, cells arrested for several hours in an anaphase-like state with condensed sister chromatids and detectable Cyclin B1 (Figure 3a, d, e, Figure 4a, d, e, Figure 6a, b, f and Videos 5 and 6).

**Figure 2.**
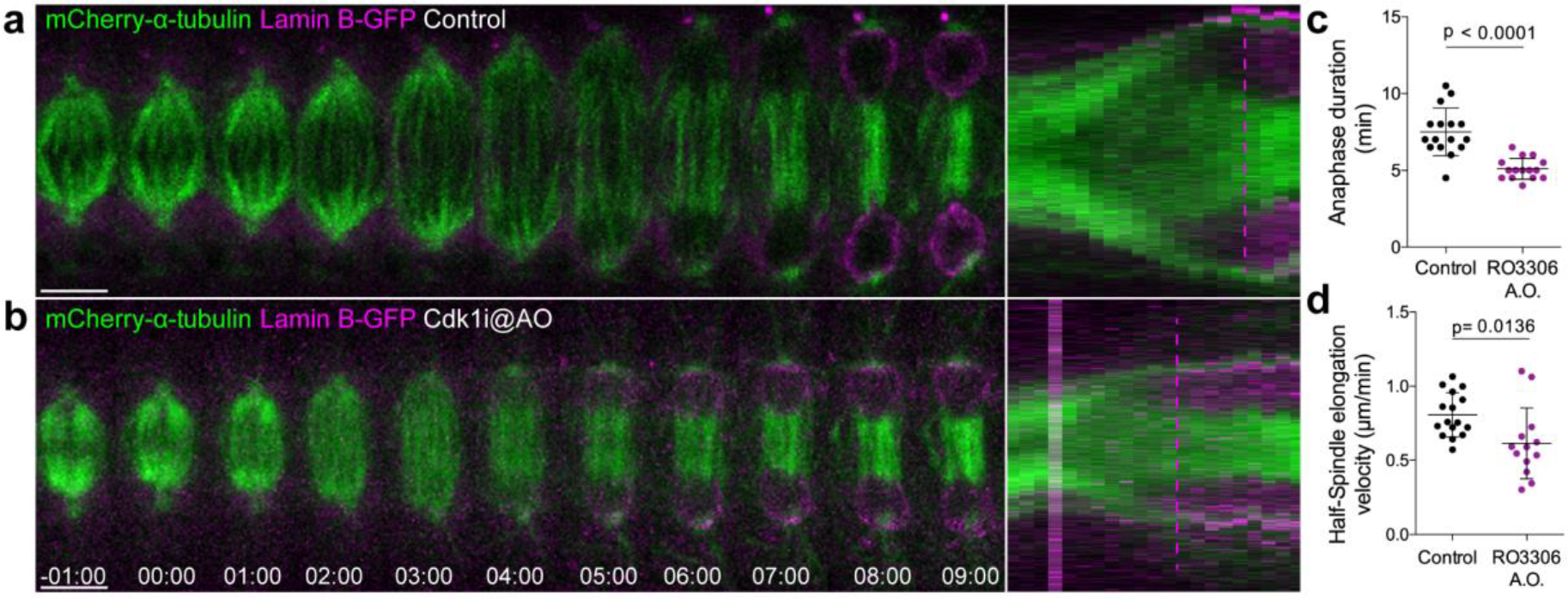
Cdk1 inhibition at anaphase onset accelerates NER. **(a)** and **(b)** Control and Cdk1-inhibited *Drosophila* S2 cells at anaphase onset (A.O.) stably expressing Lamin B-GFP/mCherry-α-tubulin. Scale bars are 5 μm. Time is in min:sec. Panels on the right side show the corresponding collapsed kymographs. Dashed lines indicate the moment of NER. **(c)** and **(d)** Quantification of anaphase duration (control n=16 cells; Cdk1i n=12 cells) and half-spindle elongation velocity (control n=16 cells; Cdk1i, n=13 cells), respectively, in the conditions shown in (a) and (b).

**Figure 3.**
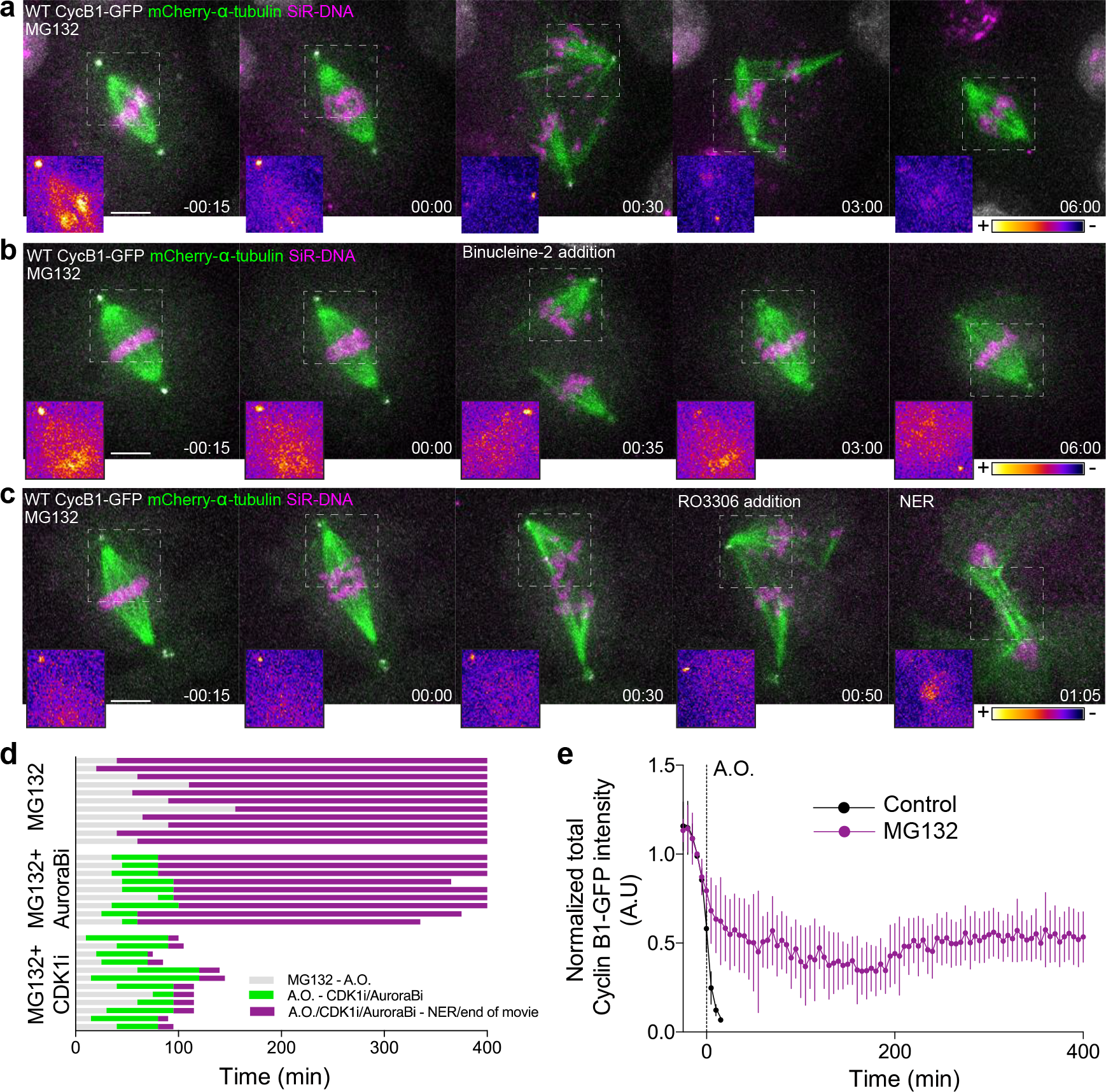
Cdk1 inactivation during anaphase licenses mitotic exit and requires proteasome-mediated degradation. **(a)** Representative *Drosophila* S2 cell stably expressing Cyclin B1-GFP/mCherry-α-tubulin and stained with SiR-DNA to follow mitotic chromosomes, showing a strong anaphase arrest after treatment with MG132 (20 μM). Importantly, cells entered anaphase with Cyclin B1 levels compared to control cells, despite the presence of MG132. **(b)** and **(c)** *Drosophila* S2 cells arrested in anaphase with MG132, and treated with Aurora B inhibitor (Binucleine-2) or Cdk1 inhibitor (RO3306), respectively, 30-60 min after the anaphase arrest. In (a), (b) and (c) a half-spindle region is highlighted with the LUT “fire” to reveal Cyclin B1-GFP fluorescence during the anaphase arrest. Scale bars in all panels are 5 μm. Time in all panels is in hh:min. **(d)** Timeline of the experiments shown in (a), (b) and (c) with quantification of total anaphase duration. A.O. = Anaphase onset. **(e)** Cyclin B1-GFP degradation profile in untreated (n=12 cells) and MG132-treated cells arrested in anaphase (n=6 cells). A.O. = Anaphase onset.

**Figure 4.**
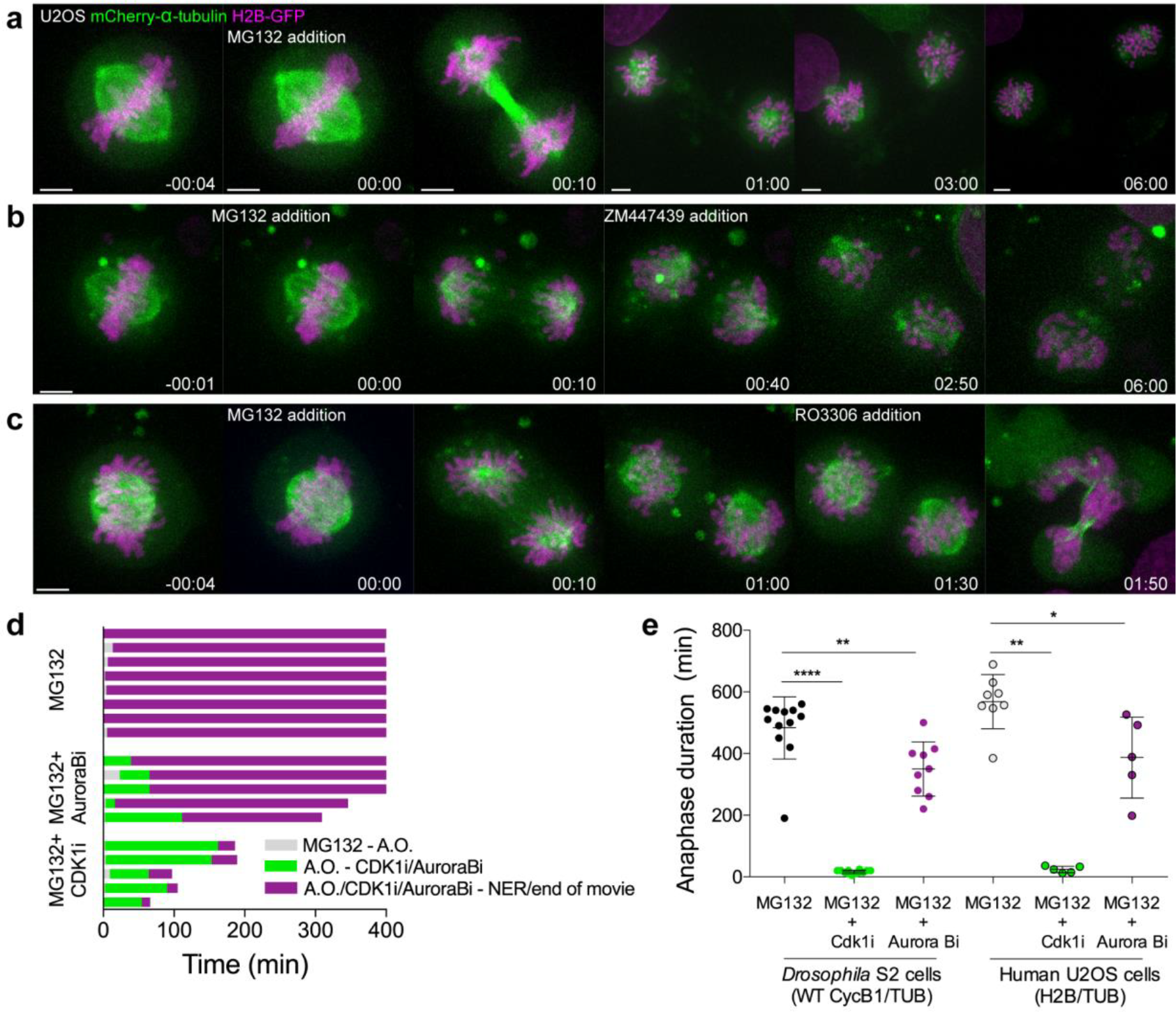
Proteasome inhibition at anaphase onset arrests human cells in anaphase in a Cdk1-dependent manner. **(a)** Representative U2OS cell stably expressing H2B-GFP/mCherry-α-tubulin treated with MG132 at anaphase onset (time 00:00). The cell became arrested in anaphase, after full chromosome separation and formation of a spindle midzone. Typically, two new spindles assembled around individual chromatids that erratically attempted to establish new “metaphase” plates. **(b)** and **(c)** Anaphase arrested U2OS cells obtained by the addition of MG132 were treated with the Aurora B inhibitor (ZM447439) or Cdk1 inhibitor (RO3306), respectively, 30-60 min after anaphase arrest. Time in all panels is in h:min. Scale bars are 5 μm. **(d)** Timeline of the experiments shown in (a), (b) and (c). **(e)** Quantification of total anaphase duration in MG132, MG132+Aurora B inhibition and MG132+Cdk1 inhibition in *Drosophila* S2 cells and human U2OS cells. Note that for MG132 and MG132+Aurora B inhibition anaphase duration corresponds to the total duration of the movie as most cells do not exit mitosis until the end of acquisition. Asterisks show significant differences (****p<0.0001; ***p<0.001; ** p<0.01 and * p<0.05).

To evaluate the respective roles of Cdk1 and Aurora B in mitotic exit, we inhibited either Cdk1 or Aurora B activity in MG132-treated anaphase cells. After Aurora B inhibition, cells remained arrested in anaphase for several hours, albeit less efficiently than controls (Figure 3b, d, Figure 4b, d, e and Videos 7 and 8). In contrast, Cdk1 inhibition immediately triggered mitotic exit (Figure 3c, d, Figure 4c-e and Videos 9 and 10). Thus, residual Cdk1 activity during anaphase is rate-limiting for mitotic exit in metazoans. Interestingly, while Aurora B inhibition was clearly not sufficient to drive cells out of mitosis if Cyclin B1 degradation was prevented, it did significantly accelerate mitotic exit under this condition, suggesting the existence of non-redundant pathways regulated by Aurora B and Cdk1.

### Cyclin B1 degradation during anaphase is mediated by APC/C^Cdh1^

APC/C^Cdc20^ regulates the metaphase-anaphase transition by targeting Cyclin B1 for degradation through recognition of a D-box, under control of the SAC (Glotzer et al., 1991). This promotes APC/C binding to Cdh1 (APC/C^Cdh1^), which recognizes substrates with either a D-box or a KEN-box (Lindon, 2008). Sequence analysis of *Drosophila* Cyclin B1 revealed a KEN box at position 248, which we mutated to AAN (Figure 5a-d). When expressed in S2 cells the KEN-Cyclin B1-GFP mutant was more slowly degraded specifically during anaphase, delaying mitotic exit (Figure 5a, b, d, f, g). These results suggest that APC/C^Cdh1^ is required for Cyclin B1 degradation during anaphase. Indeed, APC/C^Cdh1^ depletion in *Drosophila* and human cells was shown to cause an accumulation of Cyclin B1 (Ma et al., 2012; Meghini et al., 2016). To directly test this hypothesis, we performed RNAi against the *Drosophila* Cdh1 orthologue Fizzy-related (Fzr). As in the case of the KEN-box mutant, Fzr RNAi significantly delayed Cyclin B1 degradation during anaphase, postponing mitotic exit (Figure 5e-j).

**Figure 5.**
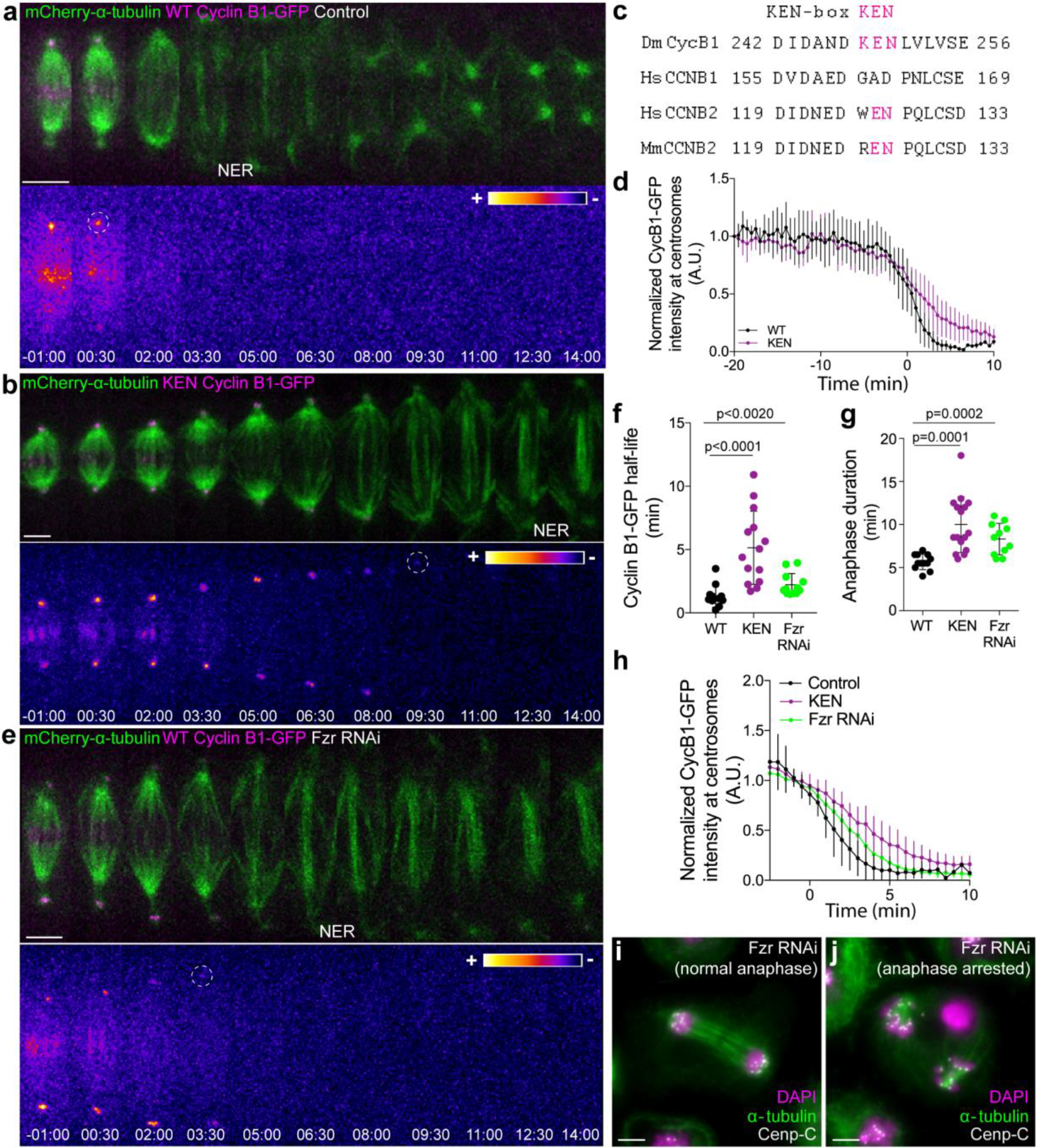
APC/C^Cdh1^ is required for Cyclin B1 degradation during anaphase and timely mitotic exit. **(a)** and **(b)** *Drosophila* S2 cells stably expressing WT Cyclin B1 or a KEN-box mutant version co-expressing mCherry-α-tubulin. **(c)** Sequence alignment showing the conservation of the *Drosophila* Cyclin B1 KEN-box with mammalian Cyclin B2, but not Cyclin B1. **(d)** Degradation profile of Cyclin B1-GFP quantified by measuring the GFP fluorescence intensity at centrosomes in control (n=6 cells) and KEN-box mutants cells (n=10 cells). Note that KEN-box Cyclin B1 degradation is only affected during anaphase. Anaphase onset = 0 min. **(e)** *Drosophila* S2 cell depleted of Fzr and expressing WT CyclinB1-GFP/mCherry-α-tubulin. Cyclin B1-GFP signal is highlighted with the LUT “fire” and dashed white circles highlight the frame before Cyclin B1 signal disappearance from centrosomes. Scale bars are 5 μm. Time in all panels is in min:sec. **(f)** and **(g)** Quantification of Cyclin B1 half-life and anaphase duration in control (n=11 cells), KEN-box mutant (n=14 cells) and Fzr-depleted cells (n=12 cells, pooled from 3 independent experiments). **(h)** Degradation profile of Cyclin B1-GFP quantified by measuring the GFP fluorescence intensity at centrosomes in control (n=12 cells), KEN-box mutants cells (n=14 cells) and Fzr-depleted cells (n=12 cells). Anaphase onset = 0 min. **(i)** and **(j)** Images of anaphase *Drosophila* S2 cells after Fzr RNAi, fixed and co-stained with Cenp-C and α-tubulin. Note that amongst apparently normal anaphase cells (i), anaphase arrested cells with clearly separated sister chromatids could also be identified (j). Scale bars are 5 μm.

Although human Cyclin B1 lacks a canonical KEN-box, its recognition by APC/C^Cdh1^ might involve a D-box or other motifs, whose inactivation compromises Cyclin B1 degradation in anaphase (Clijsters et al., 2014; Lindon, 2008; Matsusaka et al., 2014). To test the involvement of APC/C^Cdh1^ in the degradation of human Cyclin B1 during anaphase, we acutely inhibited APC/C activity at anaphase onset with Apcin and pro-TAME (Sackton et al., 2014; Zeng et al., 2010) in hTERT-RPE1 cells expressing endogenously tagged Cyclin B1-Venus. This blocked the continuous degradation of Cyclin B1 during anaphase and prevented mitotic exit for several hours, similar to proteasome inhibition (Figure 6c-f). Because pro-TAME preferentially suppresses APC/C^Cdc20^ (Zeng et al., 2010; Zhang et al., 2016) and only the simultaneous addition of both drugs at anaphase onset prevented mitotic exit (Figure 6c-f), these results suggest that a residual pool of Cyclin B1 in *Drosophila* and human cells is specifically degraded during anaphase by APC/C^Cdh1^.

**Figure 6.**
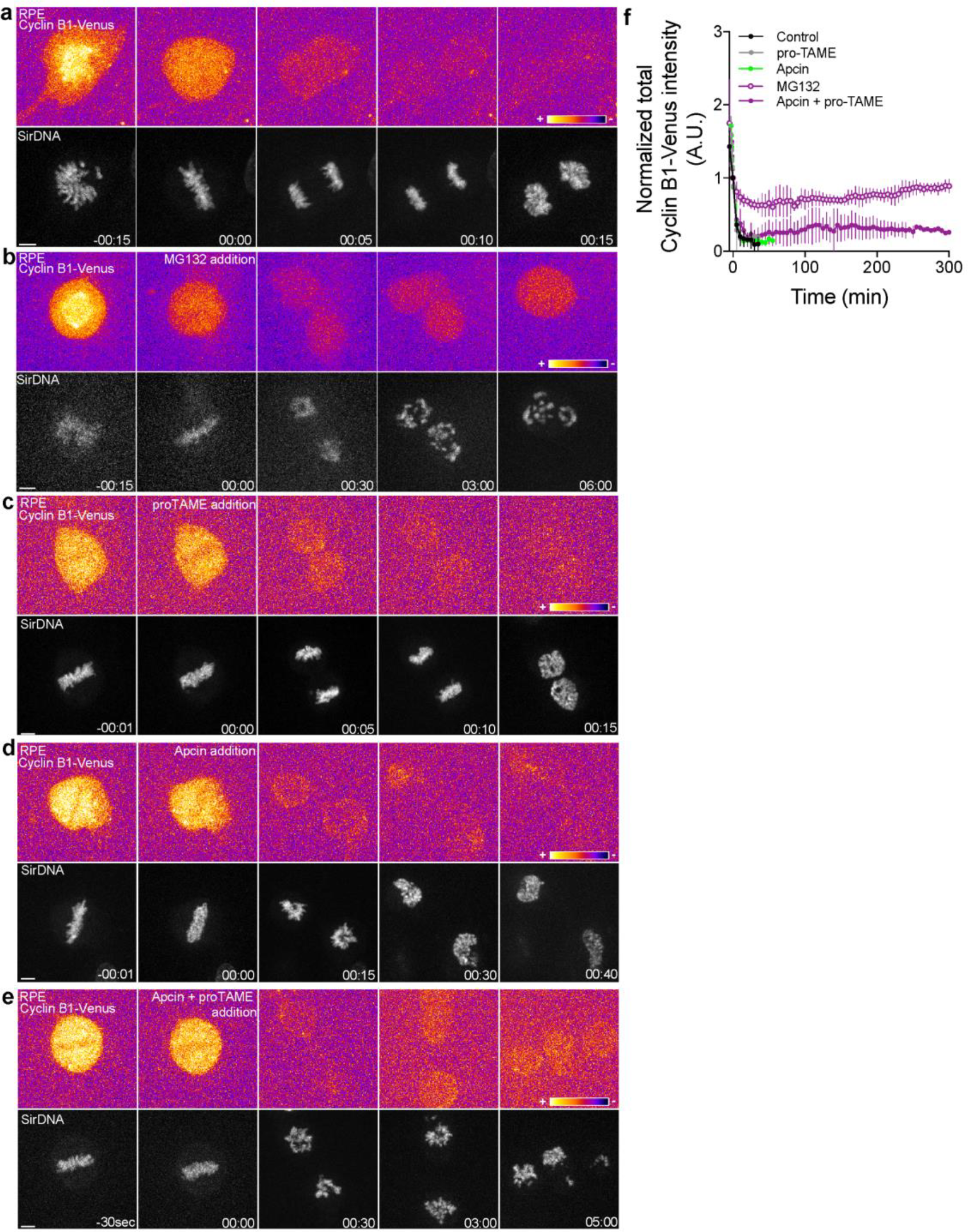
Proteasome and APC/C inhibition at anaphase onset induces an anaphase arrest in human hTERT-RPE1 cells. **(a)** Control hTERT-RPE1 cell expressing endogenous Cyclin B1-Venus and co-stained with SiR-DNA to visualize mitotic chromosomes. **(b)** MG132 addition just before or at anaphase onset (time 00:00), caused an anaphase arrest with detectable Cyclin B1 levels and condensed chromosomes. The arrest was sustained up to 6h (time window of acquisition). **(c)** pro-TAME addition at anaphase onset (time 00:00) showed no visible effect during anaphase. **(d)** Apcin addition at anaphase onset (time 00:00) induced a slight delay in DNA decondensation. **(e)** APC/C inhibition with a cocktail of pro-TAME and Apcin at anaphase onset (time 00:00) caused a strong anaphase delay (during the time window of acquisition – 5 hours). Cyclin B1 localization is highlighted with LUT “fire”. Scale bars are 5 μm. Time is in h:min. **(f)** Quantification of Cyclin B1-Venus (normalized to anaphase onset) in control (n=7 cells), MG132 (n=4 cells), pro-TAME (n=7 cells), Apcin (n=6 cells) and pro-TAME+Apcin (n=4 cells).

### Cyclin B1 localization and Cdk1 activity during anaphase depend on kinesin-6-mediated Aurora B localization at the spindle midzone

We have previously found that Aurora B inhibition abolished the cellular capacity to delay mitotic exit in response to incomplete chromosome separation during anaphase in both *Drosophila* and human cells (Afonso et al., 2014). Here we sought to investigate whether elevating Aurora B protein levels and presumably its activity impacts mitotic exit. For this purpose we overexpressed Aurora B in *Drosophila* S2 cells and found that this delayed its own transition to the spindle midzone and extended anaphase duration (Figure 7 – figure supplement 1a, b, d, e). Moreover, we found that Aurora B association with the spindle midzone is a strong predictor of anaphase duration, with several cells overexpressing Aurora B extending anaphase for more than 15 min (Figure 7 – figure supplement 1c, e, f). Interestingly, this anaphase delay depended on Cdk1 activity (Figure 7 – figure supplement 1c), suggesting that Aurora B association with the spindle midzone is able to delay mitotic exit by regulating Cdk1 activity during anaphase.

**Figure 7.**
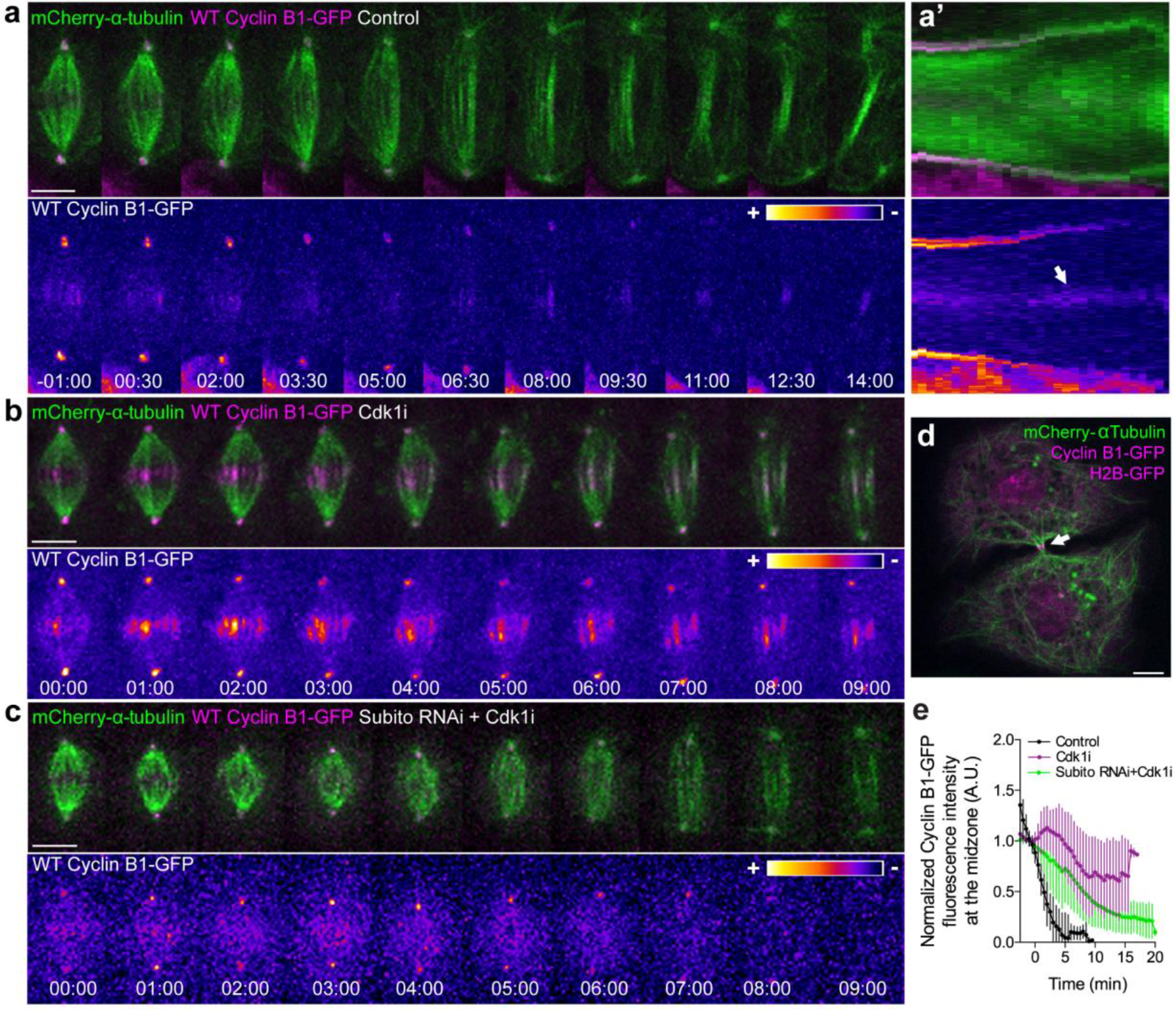
Cyclin B1 localization at the spindle midzone/midbody requires Subito/Mklp2-mediated recruitment of Aurora B. **(a)** Control *Drosophila* S2 cell stably expressing Cyclin B1-GFP/mCherry-α-tubulin showing a faint pool of Cyclin B1-GFP at the spindle midzone/midbody. **(A’)** Collapsed kymograph of the cell in a where Cyclin B1-GFP can be visualized at the spindle midzone/midbody. **(b)** *Drosophila* S2 cell treated with Cdk1 inhibitor during metaphase. Cyclin B1 is not fully degraded and becomes visibly associated with midzone microtubules as cells are forced to exit mitosis. **(c)** Cdk1 inhibition at metaphase in a *Drosophila* S2 cell expressing Cyclin B1-GFP/mCherry-α-tubulin after Subito/Mklp2 depletion by RNAi. The Cyclin B1 midzone localization is no longer detectable. Scale bars are 5 μm. Time in all panels is in min:sec. For all conditions, Cyclin B1-GFP signal is highlighted with the LUT “fire”. **(d)** Snapshot of a live *Drosophila* S2 cell expressing Cyclin B1-GFP/mCherry-α-tubulin where a midbody pool of Cyclin B1-GFP can be detected before completion of cytokinesis. Scale bar is 5 μm. **(e)** Quantification of Cyclin B1-GFP fluorescence intensity measured at the spindle midzone (identified by the mCherry-α-tubulin signal) in untreated (n=11 cells), Cdk1 inhibited cells (n=10 cells) and Cdk1 inhibition after Subito/Mklp2 depletion (n=12 cells, pooled from 2 independent experiments).

To explore this possibility we took advantage of the extremely flat morphology of *Drosophila* S2 cells growing on concanavalin-coated coverslips to investigate the precise localization of Cyclin B1 during anaphase. This allowed us to detect a faint pool of Cyclin B1-GFP associated with the spindle midzone and midbody during a normal mitosis and cytokinesis, respectively (Figure 7a, a’, d), as reported previously in *Drosophila* germ cells (Mathieu et al., 2013). Detection of the midzone pool of Cyclin B1 was enhanced by abrupt Cdk1 inactivation during metaphase, causing cells to prematurely enter anaphase with high Cyclin B1 levels (Figure 7b, e). Under these conditions, Cyclin B1 co-localized with Aurora B at the spindle midzone (Figure 7 – figure supplement 2a-c) and this localization depended on the Aurora B midzone-targeting protein Subito/kinesin-6 (Figure 7b, c, e). Altogether, these results suggest that Cyclin B1 localization and Cdk1 activity during anaphase are spatially regulated by Aurora B at the spindle midzone.

To investigate whether Aurora B at the spindle midzone regulates Cdk1 activity, we monitored Cyclin B1-GFP/Venus degradation after Subito/Mklp2/kinesin-6 RNAi in *Drosophila* S2 and human hTERT-RPE1 cells. This caused a short, but significant delay in Cyclin B1-GFP/Venus degradation during anaphase, increasing Cyclin B1 half-life and often delaying mitotic exit (Figure 8a-e, see also Figure 1 – figure supplement 3a-f). Overall, these data support a model in which the establishment of an anaphase Aurora B phosphorylation gradient concentrates Cyclin B1 and consequently Cdk1 activity at the spindle midzone.

**Figure 8.**
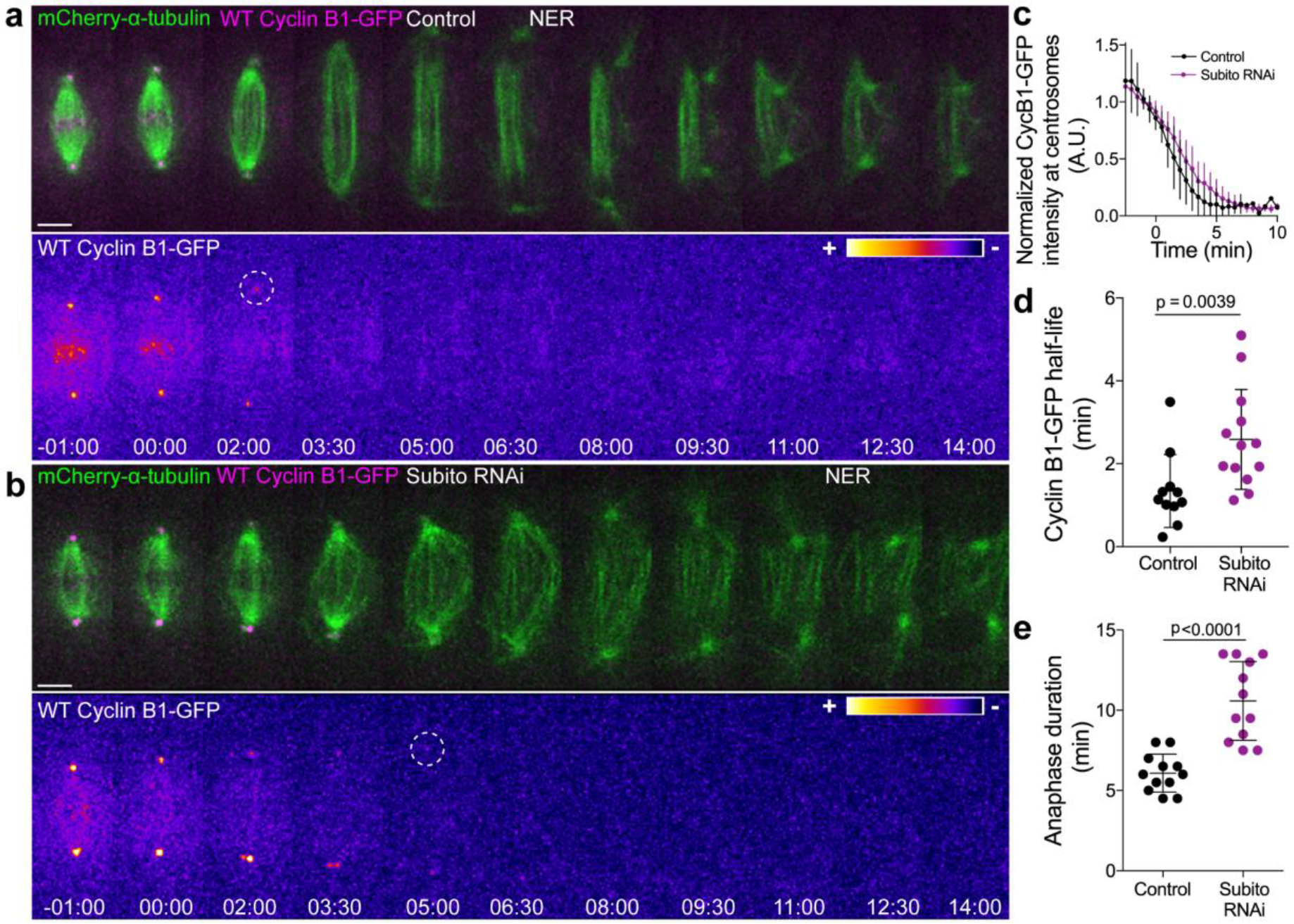
Preventing Aurora B localization at the spindle midzone delays Cyclin B1 degradation during anaphase. **(a)** and **(b)** Control and Subito/Mklp2-depleted *Drosophila* S2 cells stably expressing Cyclin B1-GFP/mCherry-α-tubulin. Cyclin B1-GFP localization is highlighted with the LUT “fire” and dashed white circles highlight the frame before Cyclin B1 signal disappearance from centrosomes. Scale bars are 5 μm. Time is in min:sec. **(c)** Degradation profile of Cyclin B1-GFP quantified by measuring fluorescence intensity at centrosomes in control (n=12 cells) and Subito/Mklp2-depleted S2 cells (n=13 cells, pooled from 3 independent experiments). **(d)** and **(e)** Calculated Cyclin B1-GFP half-life and quantified anaphase duration, respectively, in the same control and Subito/Mklp2-depleted S2 cells as in (c).

### Aurora B-mediated phosphorylation of Cyclin B1 regulates mitotic exit

Aurora B mediates Cyclin B1 phosphorylation *in vivo* and this has been implicated in the last steps of cytokinesis in *Drosophila* germ cells (Mathieu et al., 2013). As so, we investigated whether Aurora B-mediated phosphorylation affects Cyclin B1 degradation during anaphase and consequently mitotic exit. Overexpression of a non-phosphorylatable mutant (5A) in *Drosophila* S2 cells caused an average faster decay of Cyclin B1 and decreased anaphase duration compared with overexpression of wild-type (WT) or a phosphomimetic mutant (5E) (Figure 9a-e). Importantly, cells overexpressing the 5A mutant consistently entered anaphase with the lowest Cyclin B1 levels under the same induction regimens (Figure 9f), suggesting that phosphorylation of Cyclin B1 by Aurora B ensures a smooth and controlled degradation of residual Cyclin B1 during anaphase that prevents premature mitotic exit. In agreement, acute PP1/PP2A phosphatase inhibition with okadaic acid at anaphase onset did not interfere with the normal Cyclin B1 degradation profile during anaphase (Figure 9g). This also indicates that the effect of Cyclin B1 phosphorylation by Aurora B during mitotic exit is not reverted by PP1/PP2A-mediated dephosphorylation, but rather by Cyclin B1 degradation. Overall, these experiments indicate that Aurora B-mediated phosphorylation of Cyclin B1 ensures the necessary time to reach a safe chromosome separation distance before chromosome decondensation and NER, explaining how mitotic exit in metazoans is coordinated in space and time.

**Figure 9.**
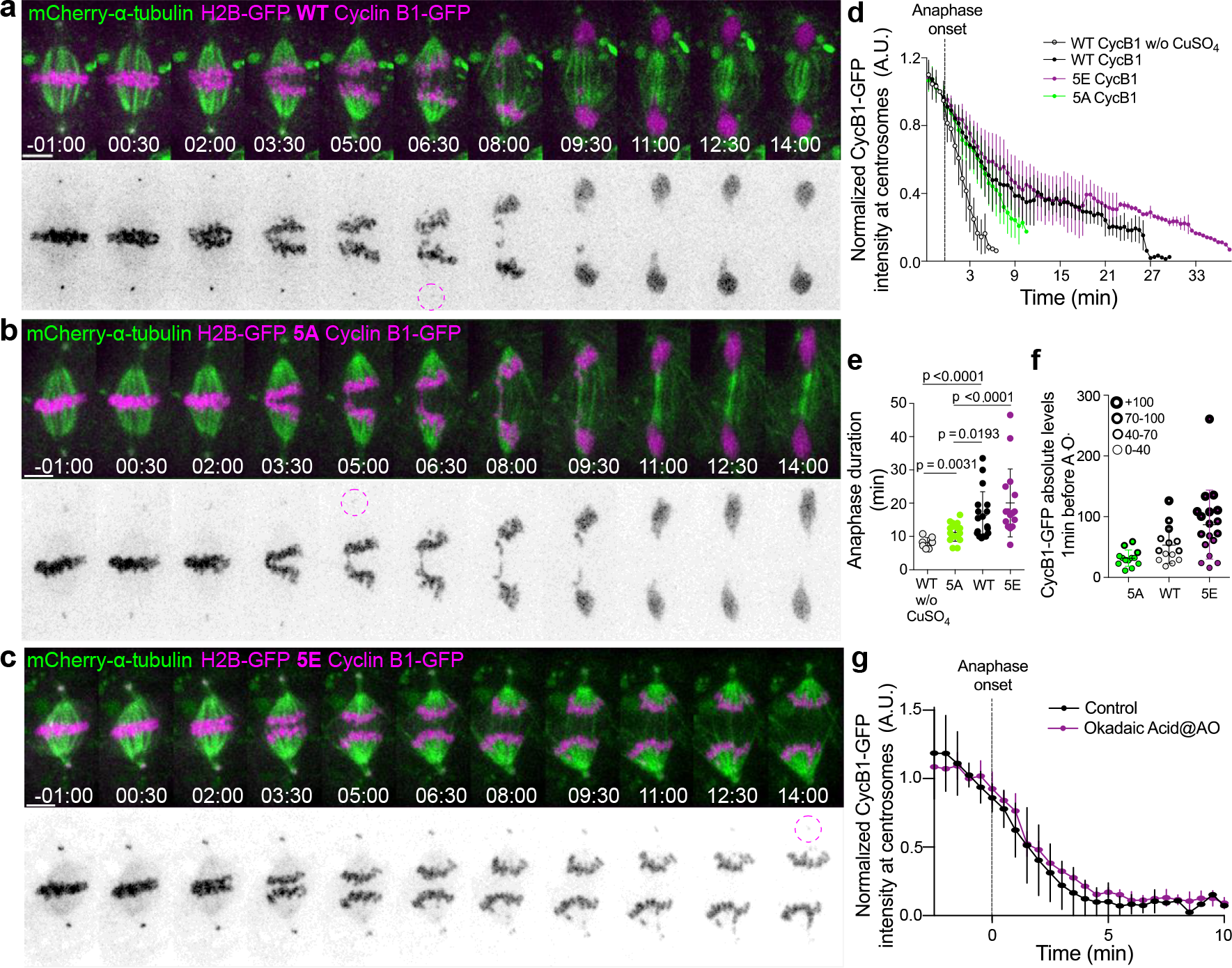
Aurora B mediated phosphorylation stabilizes Cyclin B1 and delays mitotic exit. **(a-c)** Transient overexpression of WT, non-phosphorylatable (5A) and phosphomimetic (5E) versions of Cyclin B1-GFP in a cell line stably expressing H2B-GFP/mCherry-α-tubulin. Cyclin B1-GFP signal (highlighted in inverted grayscale and with dashed magenta circle) can be detected at the centrosomes. Time in all panels is in min:sec. Scale bars are 5 µm. **(d)** Degradation profiles for the different Cyclin B1-GFP constructs as measured by fluorescence intensity at the centrosomes (WT w/o CuSO_4_=7 cells; WT=18 cells; 5A=16 cells; 5E=16 cells). **(e)** Quantification of anaphase duration in the same sample as in (d). **(f)** Absolute Cyclin B1-GFP levels with reference to 1 min before anaphase onset in WT-, 5A- and 5E-expressing cells. To facilitate data interpretation the values were distributed in four different categories revealing that 5A-expressing cells enter anaphase with the lowest Cyclin B1-GFP levels, whereas 5E-expressing cells enter anaphase with the highest. **(g)** Degradation profile of Cyclin B1-GFP measured by quantification of fluorescence intensity at centrosomes after PP1/PP2A inhibition at A.O. with okadaic acid (n=4 cells).

## Discussion

Taken together, our work reveals that Cyclin B1 degradation specifically during anaphase is rate-limiting for mitotic exit among animals that have diverged more than 900 million years ago. Most importantly, we show that Cyclin B1 homeostasis during anaphase relies on both Aurora B-mediated phosphorylation and localization at the spindle midzone. In concert with previous work (Afonso et al., 2014), our findings unveil an unexpected crosstalk between molecular “rulers” (Aurora B) and “clocks” (Cyclin B1/Cdk1) that ensures that cells only exit mitosis after proper chromosome separation, consistent with a chromosome separation checkpoint (Afonso et al., 2014; Maiato et al., 2014) (Figure 10). According to this model, APC/C^Cdc20^ mediates the abrupt Cyclin B1 degradation at the metaphase-anaphase transition under SAC control. The consequent decrease in Cdk1 activity as cells enter anaphase targets Aurora B to the spindle midzone (via Subito/Mklp2/kinesin-6); Aurora B at the spindle midzone (counteracted by PP1/PP2A phosphatases on chromatin (Vagnarelli et al., 2011)) establishes a phosphorylation gradient that spatially controls APC/C^Cdh1^-mediated degradation of residual Cyclin B1. Consequently, as chromosomes separate and move away from the spindle midzone, Cdk1 activity decreases, allowing the PP1/PP2A-mediated dephosphorylation of Cdk1 substrates (e.g. Lamin B) necessary for mitotic exit. This model is consistent with the recent demonstration that Cdk1 inactivation promotes the recruitment of PP1 phosphatase to chromosomes to locally oppose Aurora B phosphorylation (Qian et al., 2015) and recent findings in budding yeast demonstrating equivalent phosphorylation and dephosphorylation events during mitotic exit (Touati et al., 2018). Moreover, it provides an explanation for the coordinated action of two unrelated protein kinases that likely regulate multiple substrates required for mitotic exit (Afonso et al., 2017; Kettenbach et al., 2011; Petrone et al., 2016).

**Figure 10.**
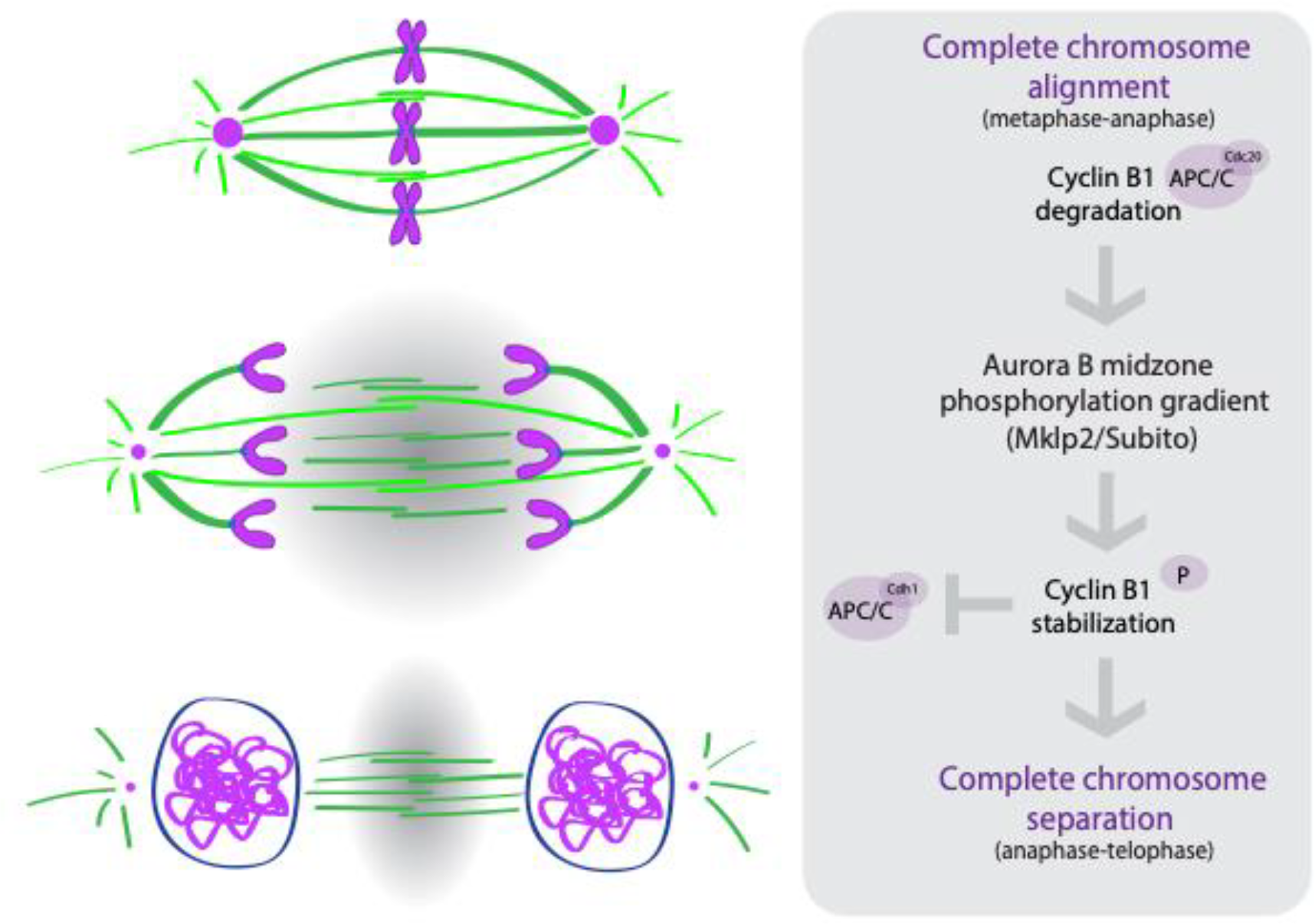
A crosstalk between molecular “rulers” (Aurora B) and “clocks” (Cdk1) licenses mitotic exit only after proper chromosome separation. Abrupt Cyclin B1 degradation at the metaphase-anaphase transition targets Aurora B to the spindle midzone (via Subito/Mklp2); Aurora B at the spindle midzone (counteracted by PP1/PP2A phosphatases) establishes a phosphorylation gradient that spatially restricts Cyclin B1 localization and delays its degradation via APC/C^Cdh1^.

Previous landmark work has carefully monitored the kinetics of Cyclin B1 degradation in living human and Ptk1 cells during mitosis, and found that Cyclin B1 was degraded by the end of metaphase, becoming essentially undetectable as cells entered anaphase (Clute and Pines, 1999). However, we noticed that, consistent with our findings, a small pool of Cyclin B1 continued to be degraded during anaphase in Ptk1 cells (Clute and Pines, 1999). Subsequent analysis with a FRET biosensor in human HeLa cells also revealed residual Cdk1 activity during anaphase (Gavet and Pines, 2010), but the significance for the control of anaphase duration and mitotic exit was not investigated in these original studies. Previous works also clearly demonstrated that forcing Cdk1 activity during anaphase through expression of non-degradable Cyclin B1 prevents chromosome decondensation and NER (Parry and O’Farrell, 2001; Wheatley et al., 1997; Wolf et al., 2006). However, while these works suggested the existence of different Cyclin B1 thresholds that regulate distinct mitotic transitions, expression of non-degradable Cyclin B1 could be interpreted as an artificial gain of function that preserves Cdk1 activity during anaphase. For example, it was shown that expression of non-degradable Cyclin B1 during anaphase “reactivates” the SAC, inhibiting APC/C^Cdc20^ (Clijsters et al., 2014; Rattani et al., 2014; Vazquez-Novelle et al., 2014). Our work demonstrates in five different experimental systems, from flies to humans, including primary tissues, that Cyclin B1/Cdk1 is rate-limiting for the control of anaphase duration and mitotic exit. Failure to degrade Cyclin B1/Cdk1 during anaphase blocked cells in an anaphase-like state with condensed chromosomes for several hours, whereas complete Cdk1 inactivation in anaphase was required to trigger chromosome decondensation and NER. Importantly, if a positive feedback loop imposed by phosphatases was sufficient to drive mitotic exit simply by reverting the effect of Cdk1 phosphorylation prior to anaphase, cells would exit mitosis regardless of the remaining Cyclin B1 pool that sustains Cdk1 activity during anaphase. The main conceptual implication of these findings is that, contrary to what was previously assumed, mitotic exit is determined during anaphase and not at the metaphase-anaphase transition under SAC control.

Our findings suggest that degradation of Cyclin B1 in anaphase is mediated, at least in part, by APC/C^Cdh1^ activity. Substrate specificity for Cdh1 is thought to be exclusively provided by the presence of a KEN box (Pfleger and Kirschner, 2000), which is present in *Drosophila* Cyclin B1. Mutation of the KEN box or Cdh1 depletion by RNAi in *Drosophila* S2 cells compromised Cyclin B1 degradation after anaphase onset, with a consequent extension of anaphase and delay in mitotic exit. Noteworthy, human Cyclin B1 lacks a canonical KEN box, but this does not imply that it cannot be degraded by APC^Cdh1^. Indeed, there are several Lysine residues in human Cyclin B1 that might compensate the lack of a canonical KEN box and might well work as a substrate recognition signal for APC/C^Cdh1^ during anaphase (Clijsters et al., 2014). Importantly, APC^Cdh1^ can still recognize Cyclin B1 through its D-box (Fang et al., 1998), but mutating the D-box in *Drosophila* or human Cyclin B1 would also interfere with its recognition by APC/C^Cdc20^ and prevent anaphase onset, precluding any subsequent analysis. In addition, because APC/C^Cdh1^ is required to degrade Cdc20 during anaphase (Pfleger and Kirschner, 2000), depleting Cdh1 from human cells might not necessarily affect mitotic exit because APC/C^Cdc20^ would still be active and able to control Cyclin B1 degradation during anaphase. For this reason, APC/C^Cdh1^ is dispensable for mitosis in mammalian cells (Garcia-Higuera et al., 2008; Sigl et al., 2009). Thus, although it is likely that APC/C^Cdh1^ mediates Cyclin B1 degradation during anaphase in both flies and humans, the most direct test to this model is the experiment provided in Drosophila cells with a Cyclin B1 KEN-box mutant. Nevertheless, we were able to show that acute APC/C inhibition with both Apcin (Sackton et al., 2014) and Pro-TAME (Zeng et al., 2010) in human hTERT-RPE1 cells expressing endogenously tagged Cyclin B1-Venus also prevented mitotic exit. Because Pro-TAME preferentially suppresses APC/C^Cdc20^ (Zhang et al., 2016) and only the simultaneous addition of both drugs at anaphase onset prevented mitotic exit, these results suggest that degradation of Cyclin B1 during anaphase in human cells is also mediated, at least in part, by APC/C^Cdh1^.

Our model also implies that residual Cyclin B1/Cdk1 in anaphase is spatially regulated by a midzone Aurora B gradient. Indeed, due to their favourable flat morphology, we were able to identify a small pool of Cyclin B1 enriched at the spindle midzone/midbody in *Drosophila* S2 cells and this localization was dependent on kinesin-6-mediated recruitment of Aurora B to the spindle midzone. Interestingly, human Cyclin B2, which contains a recognizable KEN box, localizes at the midbody and Cdk1 inactivation during late mitosis was required for the timely completion of cytokinesis in human cells (Mathieu et al., 2013). Thus, it is possible that in human cells, Cdk1 activity during anaphase is regulated not only by Cyclin B1, but also by Cyclin B2, likely mediated by APC/C^Cdh1^. This model predicts the existence of a midzone-centred Cdk1 activity gradient during anaphase, but so far, we have been unsuccessful in our attempts to target FRET reporters of Cdk1 activity (Gavet and Pines, 2010) to either microtubules or chromosomes. Future work will be necessary to test this prediction.

Finally, our experiments indicate that Aurora B-mediated phosphorylation of Cyclin B1 regulates Cyclin B1 homeostasis and consequently anaphase duration. One possibility is that Cyclin B1 phosphorylation by Aurora B spatially regulates Cyclin B1 degradation during anaphase, mediated by APC/C^Cdh1^. Because cells overexpressing non-phosphorylatable Cyclin B1 mutants enter anaphase with significantly lower levels compared to wild-type Cyclin B1 or a corresponding phosphomimetic mutant, one cannot formally exclude that this phosphorylation also affects Cyclin B1 degradation before cells have entered mitosis (via APC/C^Cdh1^) or prior to anaphase (via APC/C^Cdc20^). Additionally, Aurora B might also indirectly control Cyclin B1 during anaphase by regulating APC/C^Cdh1^ activity, as recently shown for Cdk1 (Fujimitsu et al., 2016; Zhang et al., 2016).

In conclusion, we uncovered an unexpected level of regulation at the end of mitosis and reconciled what were thought to be antagonistic models of mitotic exit relying either on molecular “clocks” or on “rulers”. These findings have profound implications to our fundamental understanding of how tissue homeostasis is regulated, perturbation of which is a hallmark of human cancers.

## Acknowledgements

We thank Juliette Mathieu, Jean-René Huynn, Eric Griffis, Claudio Sunkel, Jonathon Pines, Melina Schuh and Christian Lehner for the kind gift of reagents, and Marco Gonzalez-Gaitán for supporting O.A. during the final stages of this work. L.P.C. is the recipient of a Marie Skłodowska-Curie Action fellowship (grant agreement 746515). E.M.S. holds an FCT Investigator position and his work is supported by Fundação para a Ciência e a Tecnologia (PTDC/BEX-BCM/0432/2014). Work in the Maiato lab is supported by the European Research Council (ERC) under the European Union’s Horizon 2020 research and innovation programme (grant agreement No 681443) and FLAD Life Science 2020.

## Author Contributions

O.A. performed all the experiments in *Drosophila* and human cells in culture, L.P.C. performed the experiments in mouse oocytes, E.M.S. performed the experiments in the *Drosophila* follicular epithelium. L.T.F. established a cell line used in this study. All authors designed and analyzed their respective experiments, under the supervision of H.M. O.A. and H.M. wrote the manuscript with contributions from all the authors. H.M. conceived and coordinated the project.

---

**Figure 1 – figure supplement 1.**
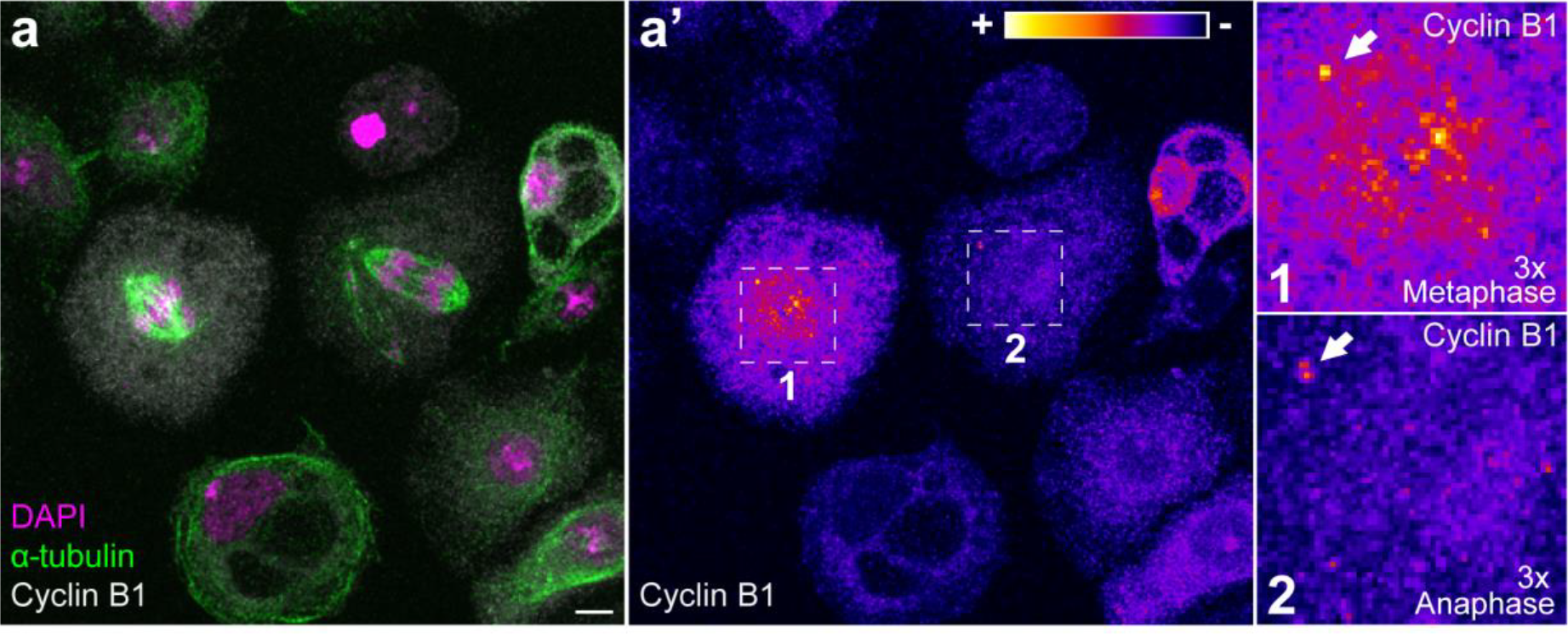
Endogenous Cyclin B1 is detectable on centrosomes in metaphase and anaphase in *Drosophila* S2 cells. **(a)** Images of fixed *Drosophila* S2 cells co-stained for endogenous Cyclin B1 and α-tubulin. **(a’)** Highlights Cyclin B1 levels with the LUT “fire”. Note the global decrease in endogenous Cyclin B1 levels from metaphase (1) to anaphase (2) (compare cytoplasmic pools). Nevertheless, early anaphase cells maintain a centrosome-associated pool of Cyclin B1 (white arrows in the magnifications). Scale bar is 5 μm.

**Figure 1 – figure supplement 2.**
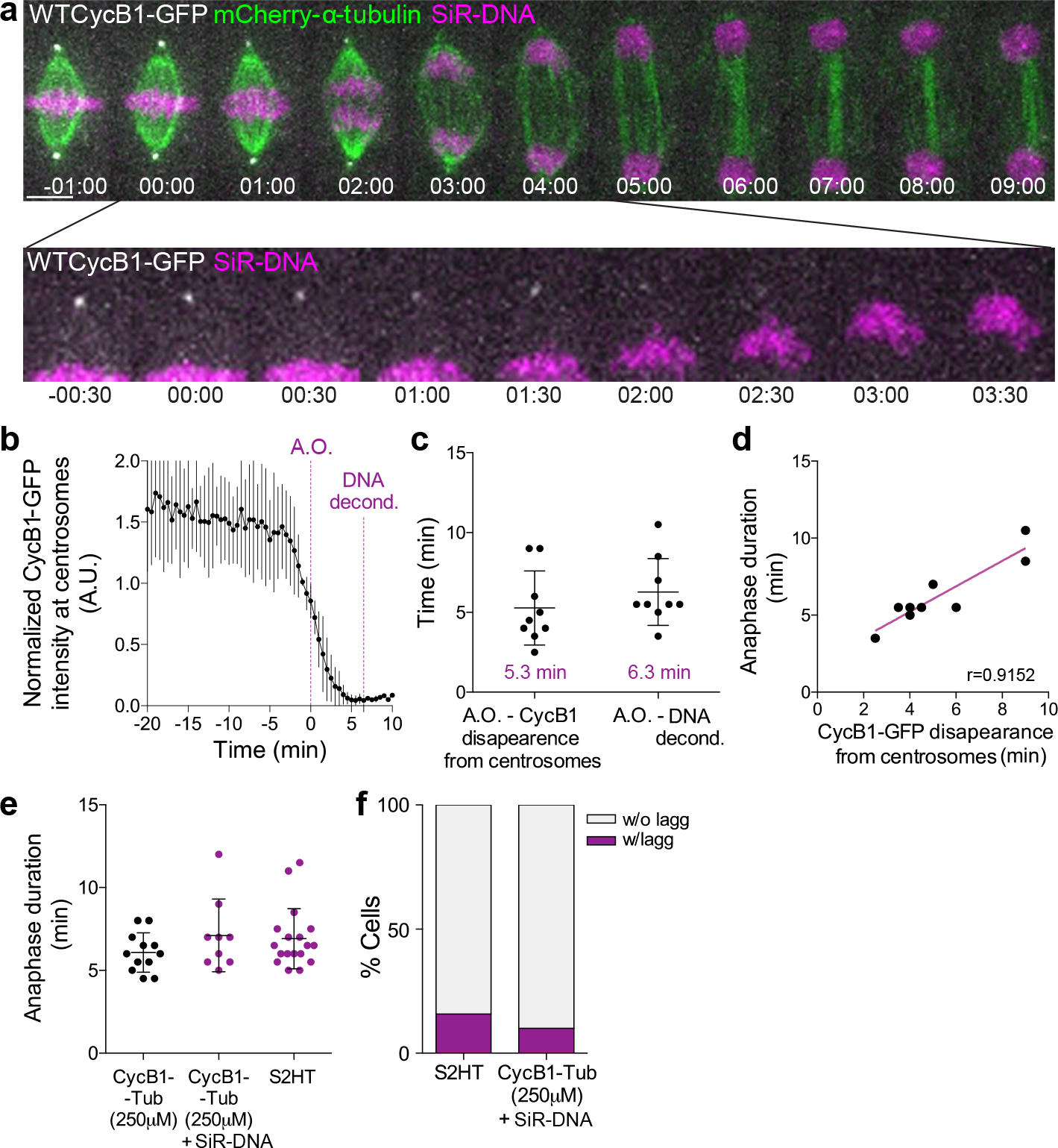
Cyclin B1 degradation during anaphase correlates with DNA decondensation. **(a)** Control *Drosophila* S2 cell stably expressing Cyclin B1-GFP/mCherry-α-tubulin and stained with SiR-DNA to follow mitotic chromosomes. Cyclin B1-GFP localization and degradation on centrosomes during anaphase is highlighted. Scale bar is 5 μm. Time is in min:sec. **(b)** Quantification of Cyclin B1-GFP fluorescence intensity at centrosomes (n=6 cells). Fluorescence intensity values were normalized to metaphase (−20 min). A.O.=Anaphase onset. **(c)** Quantification of the timing from A.O. to Cyclin B1-GFP disappearance from centrosomes and respective anaphase duration (anaphase onset to DNA decondensation) (n=9 cells). **(d)** Correlation between Cyclin B1-GFP disappearance from centrosomes and anaphase duration (p=0.0011). A parametric Pearson correlation was computed for this analysis. **(e)** Comparison of anaphase duration between *Drosophila* S2 cells expressing Cyclin B1-GFP/mCherry-α-tubulin, the same cells with SiR-DNA or cells expressing H2B-GFP/mCherry-α-tubulin. **(f)** Frequency of *Drosophila* S2 cells with lagging chromosomes in the same samples as in (b). In both (b) and (c) SiR-DNA had no impact on anaphase progression or chromosome segregation fidelity.

**Figure 1 – figure supplement 3.**
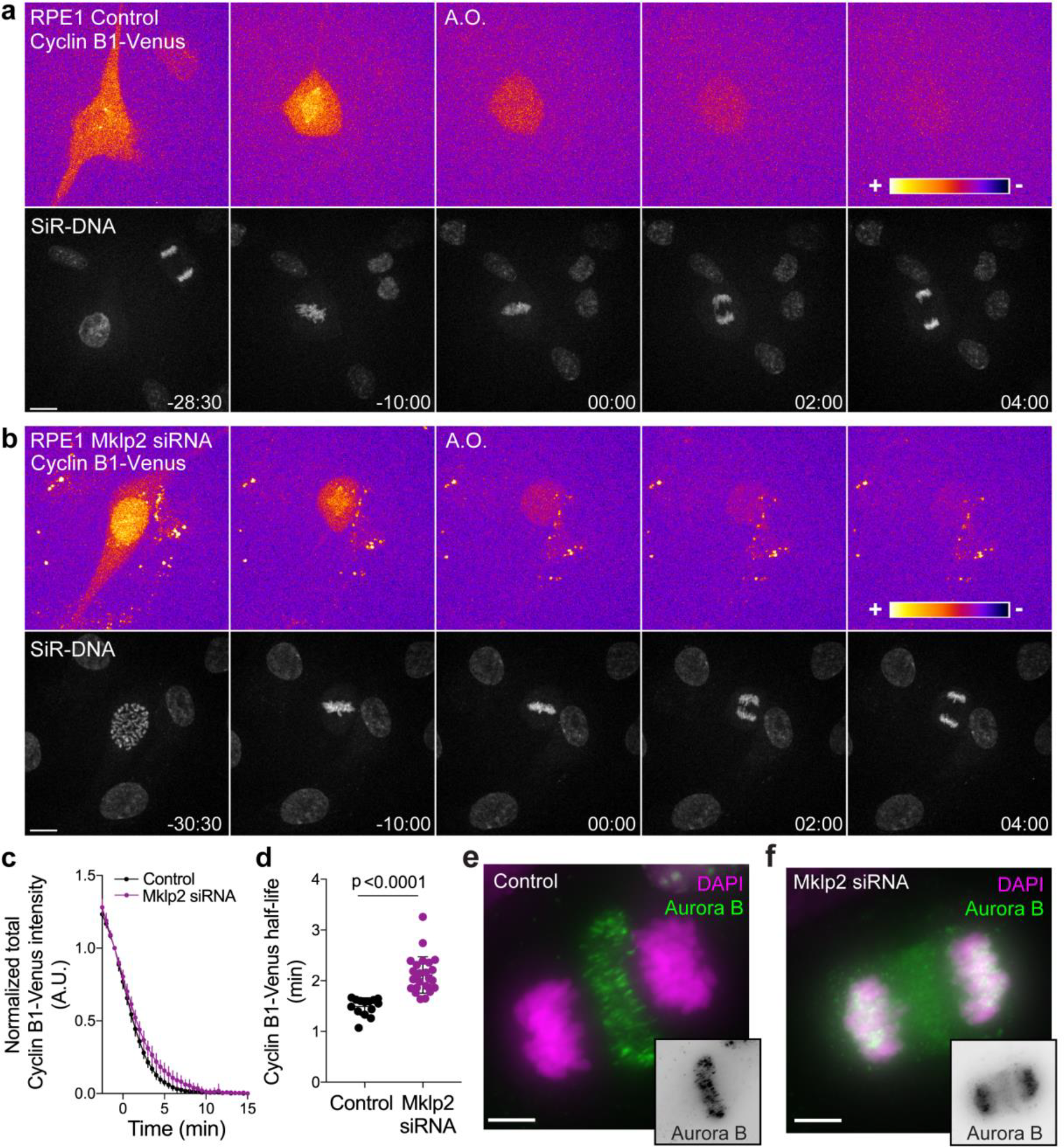
Aurora B localization at the spindle midzone impacts Cyclin B1 degradation during anaphase in hTERT-RPE1 cells. **(a)** and **(b)** Representative examples of a control and Mklp2-depleted hTERT-RPE1 cells expressing endogenous Cyclin B1-Venus and co-stained with SiR-DNA. Cyclin B1-Venus localization and levels are highlighted with the LUT “fire”, showing detectable levels of Cyclin B1 during anaphase. Time is in min:sec **(c)** Degradation profile of Cyclin B1-Venus in control (n=14 cells) and Mklp2 siRNA (n=24 cells, pooled from 2 independent experiments). Anaphase onset = 0 min. **(d)** Calculated Cyclin B1-Venus half-life after anaphase onset. **(e)** and **(f)** Aurora B staining in control and Mklp2 depleted hTERT-RPE1 cells from the same experiment shown in the live imaging in (b). Mklp2 siRNA shows a strong impairment of Aurora B translocation from chromosomes to the spindle midzone. Scale bars are 5 μm.

**Figure 2 - figure supplement 1.**
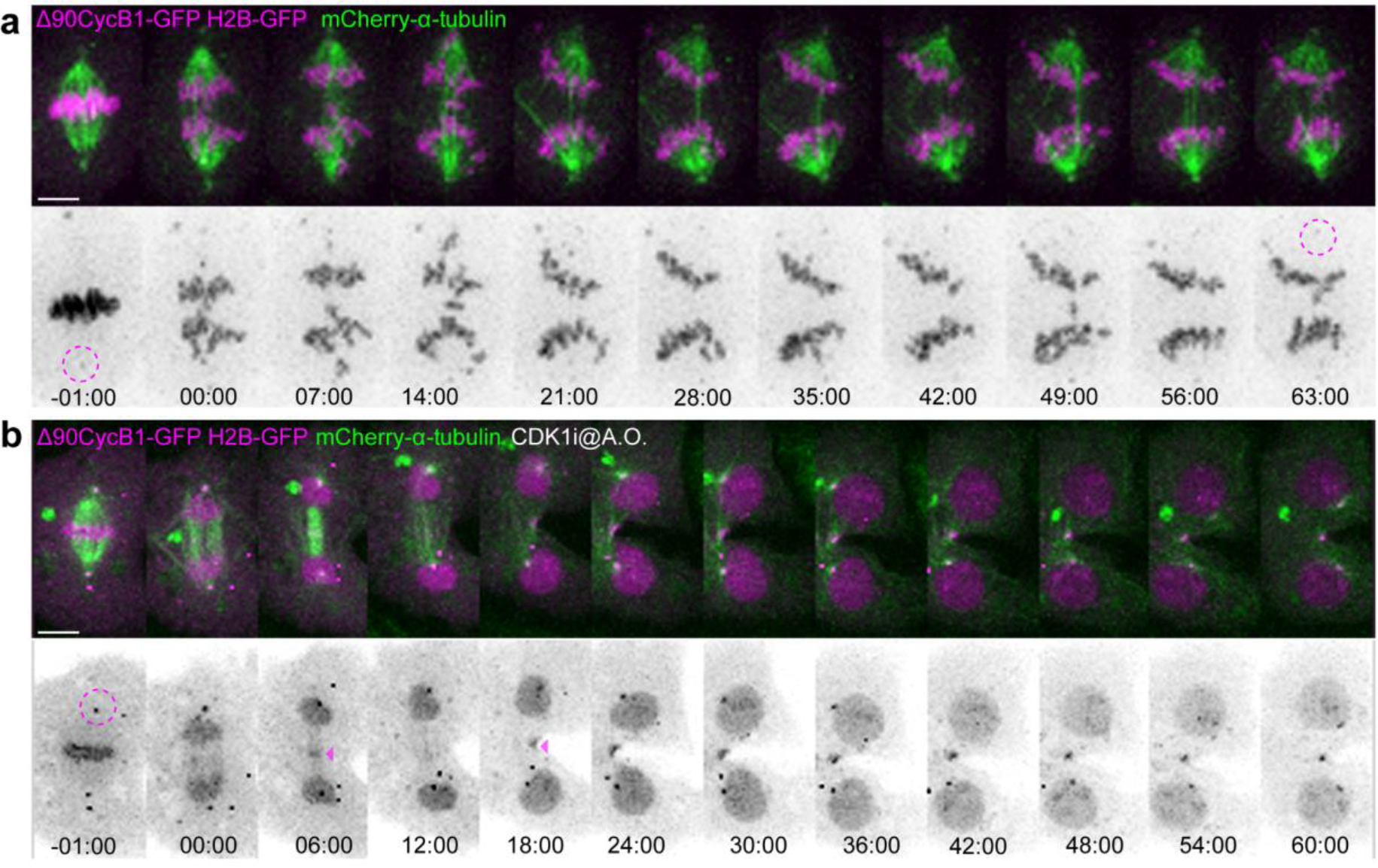
Cdk1 inhibition during anaphase is required for mitotic exit. **(a)** Control *Drosophila* S2 cell stably expressing H2B-GFP/mCherry-α-tubulin transiently expressing Δ90CycB1-GFP, showing the expected anaphase arrest. **(b)** The same experimental set-up was used for Cdk1 inhibition at anaphase onset (A.O.). Cells were forced to exit mitosis. Cyclin B1-GFP localization and H2B-GFP are highlighted in inverted grayscale. Note the centrosomal pool (magenta dashed circles) that remains during the anaphase arrest and the midzone localization of Cyclin B1-GFP after Cdk1 inhibition (magenta arrowheads). Time is in min:sec. Scale bars are 5 μm.

**Figure 7 – figure supplement 1.**
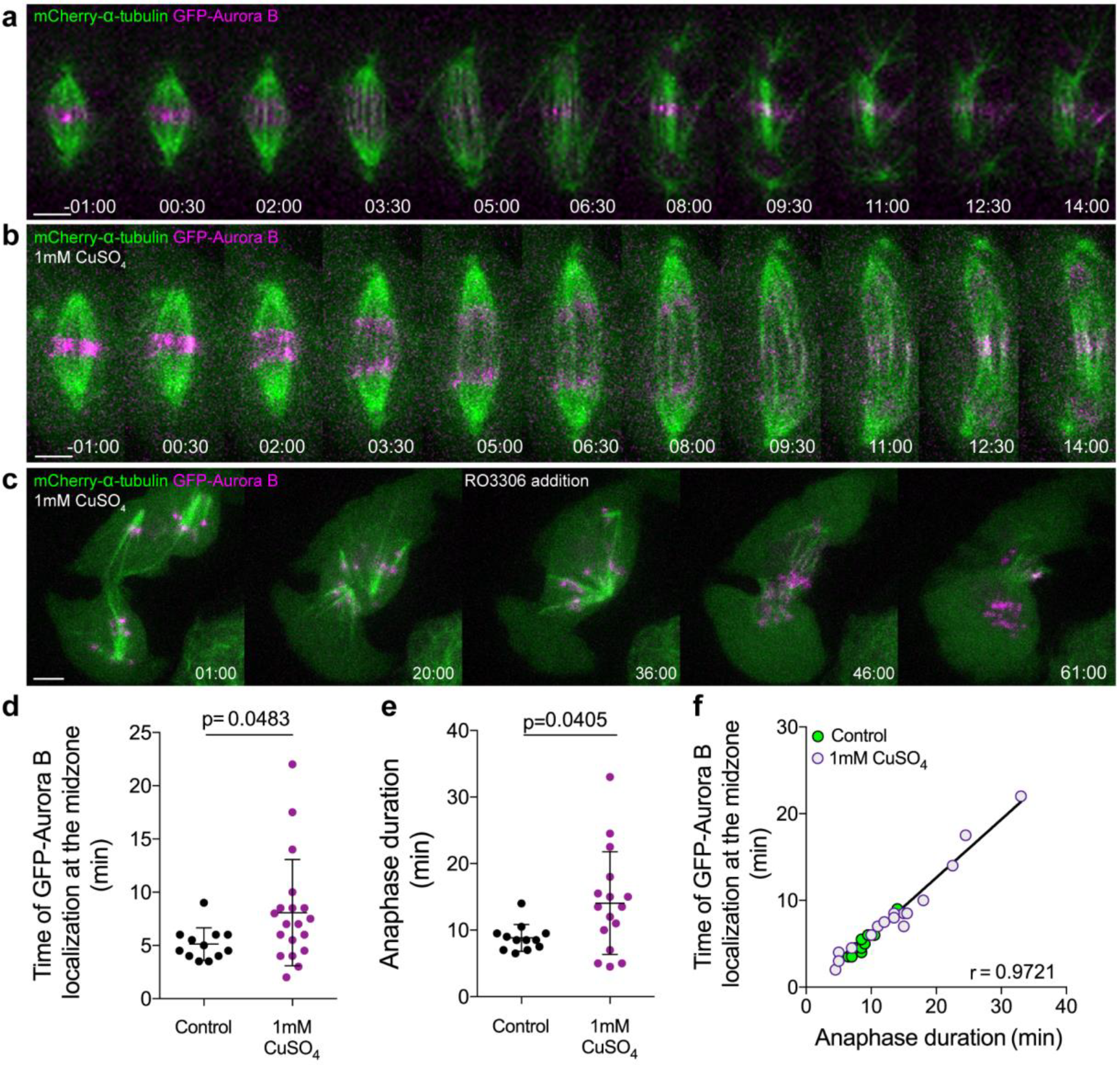
Aurora B overexpression induces a Cdk1-dependent anaphase delay. **(a)** Control *Drosophila* S2 cell stably expressing GFP-Aurora B/mCherry-α-tubulin. **(b)** *Drosophila* S2 cell overexpressing GFP-Aurora B where a delay in anaphase progression could be observed. **(c)** Cdk1 inhibition in a *Drosophila* S2 cell arrested in anaphase due to GFP-Aurora B overexpression. The cell exited mitosis immediately after drug addition. Scale bars are 5 μm. Time in all panels is in min:sec. **(d)** and **(e)** Quantification of the time of GFP-Aurora B recovery at the spindle midzone and anaphase duration, respectively, in control (n=12 cells) and GFP-Aurora B overexpressing cells (treated with 1 mM CuSO_4_) (n=16 cells). **(f)** Positive correlation (p <0.0001) between the time of Aurora B recovery at the spindle midzone and anaphase duration in control (n=12 cells) and Aurora B overexpression (n=16 cells). A nonparametric Spearman correlation was computed for this analysis.

**Figure 7 – figure supplement 2.**
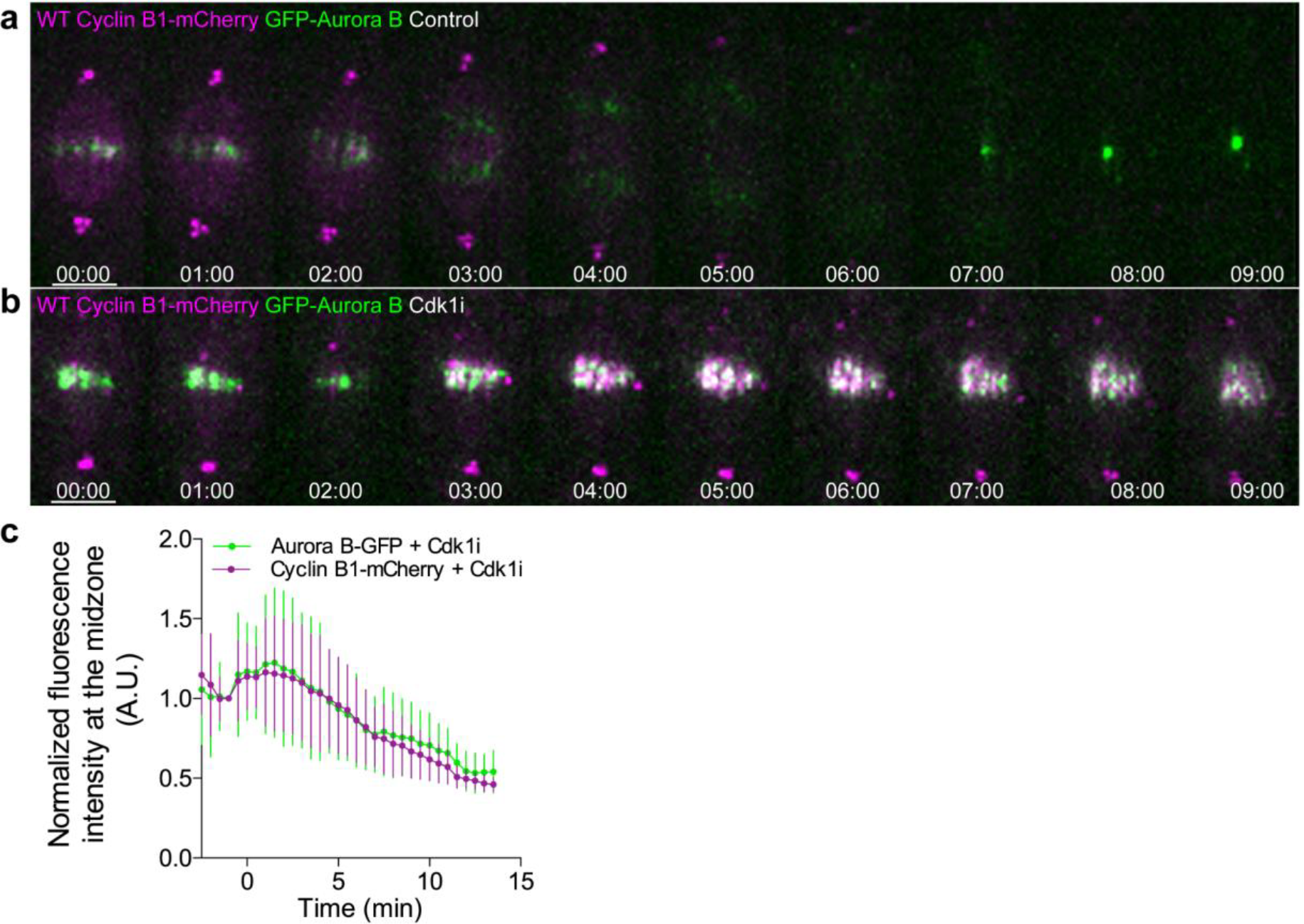
Cyclin B1 co-localization with Aurora B at the spindle midzone after Cdk1 inhibition in metaphase. **(a)** Untreated control *Drosophila* S2 cell or **(b)** treated with Cdk1 inhibitor in metaphase and expressing GFP-Aurora B/Cyclin B1-mCherry, showing a clear co-localization between both signals upon Cdk1 inhibition. Scale bars are 5 μm. Time is in min:sec. **(c)** Quantification of GFP-Aurora B and Cyclin B1-mCherry fluorescence signals at the spindle midzone after Cdk1 inhibition in metaphase (n=7 cells).

## Methods

### Cell culture

*Drosophila* S2 cells were cultured in Schneider medium supplemented with 10% FBS and grown at 25ºC. For live imaging cells were plated 2-3 h before imaging in MatTek dishes (MatTek Corporation) pre-coated with 0.25 mg/ml concanavalin A. Human U2OS and HeLa cells were cultured in DMEM supplemented with 10% FBS and grown in a 5% CO_2_ atmosphere at 37°C. hTERT-RPE1 cells were grown in DMEM/F12 medium supplemented with 10% FBS. For all human cell lines culture medium was changed to L-15 supplemented with 10% FBS 4-5 h before live imaging.

### Drug treatments

For acute inhibition of Cdk1 in S2 and U2OS cells the Cdk1 inhibitor RO3306 (Sigma) was added in metaphase or at anaphase onset, depending on the experimental set-up, at final concentration of 10 μM. Binucleine-2 (Sigma) was used to acutely inhibit Aurora B activity in *Drosophila* S2 cells at a final concentration of 40 μM in all experiments. For the live MG132 assay in *Drosophila* S2 cells, metaphase cells grown in the absence of the drug were selected in a multipoint set-up and MG132 was added after the first round of acquisition (5 min), at a final concentration of 20 μM. Cells that entered anaphase in a time window between 30-60 min after drug addition became arrested in anaphase, while the remaining cells became arrested in metaphase. MG132 addition at anaphase onset had no effect on the anaphase-telophase transition, likely due to a slow uptake of the drug in S2 cells. For the MG132 assay in U2OS and hTERT-RPE1 cells, MG132 was added 1-2 min before or at anaphase onset also at a final concentration of 20 μM. Drug addition 1-2 min after anaphase onset in U2OS or hTERT-RPE1 cells had no effect on anaphase duration. APC/C inhibition at anaphase onset on hTERT-RPE1 cells was achieved with a combination of both pro-TAME (Boston Biochem) and Apcin (Tocris Bioscience) used at final concentrations of 6.3 μM and 200 μM, respectively, as previously reported (Sackton et al., 2014). Similarly to MG132 treatment, APC/C inhibition 1-2 min after anaphase onset, or individual treatment with each drug showed a mild or no effect. ZM447439 (Tocris Bioscience) was used at 4 μM in all experiments to inhibit Aurora B in human cells. SiR-DNA (Spirochrome) was used at a final concentration of 80 nM in *Drosophila* S2 cells and 50 nM in hTERT-RPE1 cells and incubated 30-60 min prior to imaging.

### Immunofluorescence microscopy

*Drosophila* S2 cells were grown in coverslips previously pre-coated with 0.25 mg/ml concanavalin A. hTERT-RPE1 control and Mklp2-depleted cells were plated on glass coverslips. Cells were fixed with 4% PFA for 10 minutes, permeabilized with 0.3% Triton diluted in PBS for 10 min and washed 3×5 min with PBS-Tween. Primary antibodies used were mouse anti-α-tubulin (1:2000; B-512 clone, Sigma); rabbit anti-Cyclin B1 (gift from James Wakefield, University of Exeter); mouse anti-Aurora B (1:500, Aim1, BD Biosciences) and rabbit anti-Cenp C (gift from Claudio Sunkel, i3S, University of Porto, Portugal). Corresponding Alexa-Fluor secondary antibodies were used at 1:2000. Fixed images from Figure 1 – figure supplement 1 were obtained on an inverted Zeiss 780 confocal microscope with a 63× oil immersion objective. Fixed images from Figure 1 – figure supplement 3e and f were obtained on an AxioImager Z1 with a 63×, plan oil differential interference contrast objective lens, 1.4 NA (from Carl Zeiss), equipped with a charge-coupled device (CCD) camera (ORCA-R2; Hamamatsu Photonics).

### Constructs

The pMT-GFP-Aurora B and the non-degradable Cyclin B1-GFP were described previously(Afonso et al., 2014). The WT-, 5A- and 5E-Cyclin B1 pENTR constructs were a kind gift from Jean-René Huynh (Institute Curie, France). For expression in *Drosophila* S2 cells WT-, 5A- and 5E-Cyclin B1 constructs were cloned in a pMT-W-GFP destination vector (gift from Eric Griffis, University of Dundee, UK) using the LR clonase reaction from the Gateway system (Invitrogen). The KEN-box mutant was generated in the pMT-Cyclin B1-GFP plasmid using a site-directed mutagenesis kit (Agilent), following the manufacturer instructions. The following primers were used for mutagenesis: Fw: 5’-GGACATTGATGCCAATGACGCGGCGAACCTGGTACTGGTCTCC-3’ and Rv 5’-GGAGACCAGTACCAGGTTCGCCGCGTCATTGGCATCAATGTCC-3’.

### Cell lines, transfections and RNAi

The H2B-GFP/mCherry-α-tubulin and Lamin B-GFP/mCherry-α-tubulin stable cell lines have been previously described (Afonso et al., 2014). The WT-Cyclin B1-GFP/mCherry-α-tubulin, KEN-Cyclin B1-GFP/mCherry-α-tubulin and GFP-Aurora B/WT-Cyclin B1-mCherry stable cell lines were established using Effectene (Qiagen) according to the manufacturer instructions. For visualization of WT-Cyclin B1-GFP, WT-Cyclin B1–mCherry and GFP-Aurora B signals 250 μM of CuSO_4_ was added to the medium overnight (8-12 h) to induce expression from a metallothionein promoter. With this copper concentration, we obtained a mild expression of the constructs without any detectable impact on mitotic progression. Aurora B overexpression was achieved by adding 1 mM of CuSO_4_ instead of 250 μM for the same incubation period. For the analysis of the Cyclin B1 phosphorylation mutants, transient transfections on a H2B-GFP/mCherry-α-tubulin stable cell line were performed using Effectene (Quiagen) according to the manufacturer instructions and overexpression was induced with 500 μM of CuSO_4_ incubated overnight in the cell culture medium. The HeLa and hTERT-RPE1 Cyclin B1-Venus cell lines (Collin et al., 2013) were a kind gift from Jonathon Pines (Institute of Cancer Research, UK). The H2B-mRFP expression construct for human cells was obtained from Addgene (#26001), transfected to 293T cells to produce viruses and infect HeLa Cyclin B1-Venus expressing cells to generate a stable cell line. The H2B-GFP/mCherry-α-tubulin U2OS cell line and the Subito/Mklp2 RNAi in *Drosophila* S2 cells were previously described(Afonso et al., 2014). Fzr RNAi in *Drosophila* S2 cells was achieved after two rounds of RNAi treatment for a total of 7 days. The following primers were used for dsRNA synthesis: 5’-TAATACGACTCACTATAGGGCACCGGATAATCAATACTTGGC-3’ and 5’-TAATACGACTCACTATAGGGATTCAGAACGGACTTGTTCTCC-3’. Mklp2 depletion in hTERT-RPE1 cells was achieved 72 h after transfection of siRNA oligonucleotides using Lipofectamine RNAiMAX (Invitrogen) following the manufacturer instructions. The oligonucleotide sequence used was: 5’-AACGAACUGCUUUAUGACCUA-3’.

### Time-lapse microscopy

Live imaging data in figures 2 and 9 and figure 2 - figure supplement 1 were obtained from a spinning disc confocal system (Andor Technology, South Windsor, CT) equipped with an electron multiplying CCD iXonEM+ camera and a Yokogawa CSU-22 U (Yokogawa Electric, Tokyo) unit based on an Olympus IX81 inverted microscope (Melville, NY). Two laser lines (488 and 561 nm) were used for near simultaneous excitation of GFP and mCherry/mRFP and the system was driven by Andor IQ software. Time-lapse image stacks of 0.8 μm steps were collected every 30 sec, with a Plan-APO 100×/1.4 NA oil objective, with the exception of the experiments in figure 2 - figure supplement 1 where image stacks were obtained every 60 sec. Live imaging data in figures 1, 3, 4, 5a-h, 6-8, figure 1 – figure supplement 2 and 3a-d and figure 7 – figure supplement 1 and 2 were acquired in a temperature-controlled Nikon TE2000 or a Ti microscope, both equipped with a Yokogawa CSU-X1 spinning-disc head, imaged on an Andor iXon+ DU-897E EM-CCD. Excitation comprises three lasers (488nm, 561nm and 647nm) that were shuttered by an acousto-optic tunable filter (TE2000) or electronically (Ti). Sample position was controlled via a SCAN-IM Marzhauser stage and a Physik Instrumente 541.ZSL piezo (TE2000) or via a Prior Scientific ProScan stage (Ti). Imaging of *Drosophila* S2 cells was performed with a 100× Plan-Apo DIC CFI Nikon objective in all experiments. Imaging of human cells and the *Drosophila* follicular epithelium was done with an oil-immersion 60× 1.4 NA Plan-Apo DIC CFI (Nikon, VC series). Imaging of the mouse oocytes was performed either with a 60× 1.27 NA CFI Plan-Apo IR water-immersion objective (Nikon) or a 40× 1.30 NA Plan-Fluor oil-immersion objective (Nikon). Images were acquired with a 1 µm z-stack and 30 sec time-lapse interval for all live imaging experiments in *Drosophila* S2 cells, *Drosophila* follicular epithelium and human cells. Mouse oocytes were acquired with a 4 µm z-stack and a 2 min time interval. Data from figures 3 and 6 were obtained with a 5 min time interval and data from figure 4 were obtained with a 1min time interval. A temperature-controlled chamber was set up to 25ºC for *Drosophila* live imaging or to 37ºC for live imaging of human cells and mouse oocytes. SiR-DNA (Spirochrome) was used at a final concentration of 80 nM in *Drosophila* S2 cells or 50 nM in hTERT-RPE1 cells and incubated 30-60 min prior to imaging.

### *Drosophila* strains and egg chamber preparation

Flies homozygous for the eGFP protein trap insertion into *CycB1* loci (CC01846(Buszczak et al., 2007)) and expressing His2Av-mRFP (genotype: *w*; *P{PTT-GC}CycB*^*CC01846*^; *P{His2Av-mRFP}*/+) were generated to image Cyclin B1 expression at endogenous levels while monitoring mitotic progression. *Drosophila* egg chambers were dissected and mounted in the gas-permeable oil 10S VOLTALEF^®^ (VWR chemicals) for live imaging.

### *In vitro* culture and micro-injection of mouse oocytes

All mice were maintained in a specific pathogen-free environment according to the Portuguese animal welfare authority (Direcção Geral de Alimentação e Veterinária) regulations and the guidelines of the Instituto de Investigação e Inovação em Saúde animal facility. Oocytes were isolated from ovaries of 7-10 week old CD-1 mice, cultured in M2 medium under mineral oil and matured to metaphase II in vitro for 16 hours at 37ºC. Oocytes were then microinjected with mRNA encoding Cyclin B1-mCherry using an electric-assisted microinjection system (FitzHarris et al., 2018). A function generator (GW Instek AFG-2005) was used to produce electrical current instead of an intracellular electrometer. Oocytes were then transferred to M2 medium supplemented with SiR-DNA (500 nM, Spirochrome) and allowed to express the mRNA for 4-6 hours. Oocytes were then induced parthenogenically by rinsing into calcium-free M2 medium with 10 mM strontium chloride, and imaged.

### mRNA *in vitro* transcription

pGEMHE-CyclinB1-mCherry was a kind gift from Melina Schuh (Max-Planck Institute for Biophysical Chemistry). The plasmid was linearised, and capped mRNA was synthesised using the T7 ARCA mRNA kit (New England Biolabs) according to the user’s manual and resuspended in water. The final concentration of mRNA in the injection needle was 400 ng/µl.

### Definition of mitotic exit

Mitotic exit was defined either by the moment of NER or chromosome decondensation. When a nuclear envelope marker was present (ex: Lamin B in figure 2) mitotic exit was defined as the first frame of appearance of Lamin B in the nascent nucleus. In experiments where cells expressed mCherry-α-tubulin, NER was determined based on the first frame where soluble mCherry-α-tubulin was excluded from the main nucleus. In those cases where mCherry-α-tubulin was not present, SiR-DNA was used instead and visual inspection of DNA decondensation was used to define mitotic exit.

### Quantification of Cyclin B1 decay during anaphase

Image processing and quantification was performed using Fiji. Data from *Drosophila* S2 in figures 3e, 5d and h, 8c, 9d and g and, figure 1 – figure supplement 2b, Cyclin B1 were measured with a constant circular ROI around the centrosomes. This resulted in more robust measurements, which were less prone to fluorescence fluctuations in the cytoplasm. This was essential to detect differences in the Cyclin B1 decay in the different conditions tested. Exceptionally, and to compare Cyclin B1 levels in *Drosophila* and human cells, in figure 1c total Cyclin B1-GFP fluorescence intensity was measured using an ROI defined by the limits of the cell. Cyclin B1 levels from figure 7e and figure 7 – figure supplement 2 were measured with a constant rectangular ROI defined by the size of the metaphase plate before cells entered anaphase. This size and position was maintained to measure the intensity of Cyclin B1 that localized at the spindle midzone in control cells and after Cdk1 addition. Note that with the same ROI in figure 7 – figure supplement 2 both Cyclin B1 and Aurora B intensities were measured. Data from human cells from figures 1b, 6f and Figure 1 – figure supplement 3, total Cyclin B1 were measured with an ROI defined by the limits of the cell. In both cases, the ROI was used to manually track the centrosome/midzone/cell over time. Images were maximum intensity projected and background was measured for all time points with the same ROI used for Cyclin B1 measurements in a region outside the cell. The background levels were subtracted from the absolute intensity values. In the *Drosophila* follicular epithelium Cyclin B1 levels were measured in sum-intensity Z projections, using a circular ROI that was manually tracked in an homogeneous cytoplasmic region (free of accumulated Cyclin B at microtubules or centrosomes) of dividing cells. Background was measured with the equivalent ROI in G1 phase cells (identified by undetectable Cyclin B1-GFP expression) within the same egg chamber, and subtracted to dividing cells to establish the baseline background value. In mouse oocytes, Cyclin B1 was measured using a ROI placed manually into the center of the oocyte, background levels were measured in a ROI outside the cell and used for subtraction from the mean cytoplasmic intensity. Images were sum-intensity Z projected.

### Calculation of Cyclin B1 half-life

The half-life of Cyclin B1-GFP or Cyclin B1-Venus was calculated based on the formula 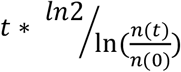 where *t* is the time interval, *n(0)* is the background subtracted Cyclin B1-GFP fluorescent intensity at anaphase onset and n(t) is the background subtracted Cyclin B1-GFP fluorescent intensity at time 4.5 min after anaphase onset.

### Statistical analysis

Normality of the samples was determined with a D’Agostino & Pearson test. Statistical analysis for two-sample comparison, with normal or non-normal distribution, was performed with a t-test or Mann-Whitney test, respectively. P value was considered extremely significant if p<0.0001 or p<0.001, respectively, very significant if 0.001<P<0.01 and significant if 0.01<P<0.05. In all plots error bars represent standard deviation. All statistical analysis was performed with GraphPad Prism V7 (GraphPad Software, Inc.).

### Code Availability

Matlab based custom routines were used for the generation of kymographs. Code is available from the corresponding author upon request.

### Data availability

All data generated or analysed during this study are included in this published article (and its supplementary information files). Raw movies are available from the corresponding author on reasonable request.

